# Proliferation in malaria parasites: how resource limitation can prevent evolution of greater virulence

**DOI:** 10.1101/2023.12.16.571859

**Authors:** Damie Pak, Tsukushi Kamiya, Megan A. Greischar

## Abstract

For parasites, robust proliferation within hosts is crucial for establishing the infection and creating opportunities for onward transmission. While faster proliferation enhances transmission rates, it is often assumed to curtail transmission duration by killing the host (virulence), a tradeoff constraining parasite evolution. Yet in many diseases, including malaria, the preponderance of infections with mild or absent symptoms suggests that host mortality is not a sufficient constraint, raising the question of what restrains evolution towards faster proliferation. In malaria infections, the maximum rate of proliferation is determined by the burst size, the number of daughter parasites produced per infected red blood cell. Larger burst sizes should expand the pool of infected red blood cells that can be used to produce the specialized transmission forms needed to infect mosquitoes. We use a within-host model parameterized for rodent malaria parasites (*Plasmodium chabaudi*) to project the transmission consequences of burst size, focusing on initial acute infection where resource limitation and risk of host mortality are greatest. We find that resource limitation restricts evolution towards higher burst sizes below the level predicted by host mortality alone. Our results suggest resource limitation could represent a more general constraint than virulence-transmission tradeoffs, preventing evolution towards faster proliferation.

## Introduction

Faster proliferation helps parasites establish infections, persist through the onslaught of host immunity, and transmit at higher rates. At the same time, aggressive proliferation can increase virulence (infection-induced host mortality), so that parasite fitness is thought to be maximized at an intermediate—rather than maximal—rate of proliferation (e.g., Anderson & May, 1982; Ewald, 1983; Frank, 1996). One key example of support for a transmission-virulence tradeoff comes from HIV where greater replicative capacity increases viral load (Prince *et al*., 2012), increasing infectiousness but accelerating progression to AIDS and eventual death (Shirreff *et al*., 2011; Fraser *et al*., 2014). An intermediate viral load maximizes cumulative transmission potential, infectiousness integrated over the duration of infection (Fraser *et al*., 2014). Yet even if host mortality is costly to transmission, low case fatality rates (0.001 to 0.01%) reported in many diseases raise the question of whether transmission-virulence tradeoffs can serve as a general explanation for lack of evolution towards ever greater rates of proliferation (reviewed in Bull & Lauring, 2014). It remains unclear if the proliferation rates of pathogenic organisms are limited primarily by the cost of host mortality, or if not, what other mechanisms are involved.

The biology of malaria parasites (*Plasmodium spp.*) conforms to key assumptions required for a transmission-virulence tradeoff, since the cycle of proliferation within the host requires destruction of red blood cells (RBCs) whose role in oxygen transport renders them crucial for host survival. Malaria parasites infect and multiply within host RBCs, which subsequently rupture to release daughter parasites (merozoites) that can invade further RBCs to continue the proliferative cycle. The upper limit on proliferation rates is determined by the burst size, the number of merozoites emerging from an infected RBC upon bursting. Parasites are rarely expected to attain maximal rates of proliferation, due in part to the need to allocate some fraction of invaded RBCs to the production of gametocytes (“transmission investment”), the specialized forms capable of infecting mosquito vectors. Faster proliferation is predicted to improve transmission success because it generates a large pool of invaded RBCs that can subsequently be committed to the production of gametocytes (Koella & Antia, 1995; Mideo & Day, 2008; Greischar *et al*., 2016, 2019). Rapid proliferation also allows malaria parasites to establish infections more successfully by overwhelming the host’s innate immune response (Mackinnon *et al*., 2002; Metcalf *et al*., 2011). However, infection can end abruptly due to catastrophic depletion in RBCs, leading to host death (Lamikanra *et al*., 2007). Accordingly, highly transmissible strains cause greater depletion in RBCs which can lead to severe symptoms like catastrophic anemia (Mackinnon & Read, 1999, 2004). Experimental rodent malaria *Plasmodium chabaudi* infections suggest that killing the host significantly reduces parasite transmission (Mackinnon *et al*., 2002). Hence, parasite traits that enable faster proliferation—including larger burst sizes and restrained transmission investment—would be expected to carry both a fitness benefit and a cost.

In addition to fitness consequences, the traits underpinning proliferation rates can only evolve if there is heritable variation on which natural selection can act. Transmission investment varies across and even within species (reviewed in Taylor & Read, 1997) and is known to be evolvable, due to the reduction and even loss of the ability to produce gametocytes in human malaria strains maintained in artificial culture (reviewed in Bousema & Drakeley, 2011). Burst sizes also vary considerably within and across species, suggesting potentially heritable variation. Human malaria (*P. falciparum*) strains differ in their maximal and median burst sizes (Reilly *et al*., 2007), and the rodent malaria species *P. chabaudi* and *P. berghei* exhibit burst size ranges of 2-12 and 6-20 merozoites/infected RBC, respectively (Garnham *et al*., 1966). Burst size even appears to vary over the course of infection (Mideo *et al*., 2011). Nonetheless, it remains an open question how selection has shaped variation in burst sizes and whether existing variation could be adaptive (reviewed in Mideo & Reece, 2012).

One key issue is whether malaria-induced host mortality is common enough to influence the evolution of burst sizes. Most human malaria cases are subclinical without severe symptoms (Lindblade *et al*., 2013; Huang *et al*., 2017). Parasite abundance only increases the probability of severe outcomes beyond a threshold, with many human infections falling below that threshold (Cunnington *et al*., 2013). Further, while larger parasite loads increase risk of mortality in human infections, greater abundance also extends the time until clearance (Mackinnon & Read, 2004). Since faster proliferation extends the time until host recovery, and host mortality may be too rare to impose an important selection pressure, other mechanisms may be needed to restrain evolution towards larger burst sizes. Although originally framed at the epidemiological rather than within-host scale, theory suggests the potential for tradeoffs between invasion and persistence (King *et al*., 2009), wherein strains that invade host populations more efficiently may deplete susceptible hosts so quickly that they drive themselves extinct. Likewise, parasites that proliferate too quickly may rapidly deplete host resources (for malaria parasites, RBCs) and thereby hinder subsequent proliferation.

Disentangling the processes that could restrict the evolution of burst size requires careful consideration of the timing of fitness costs and benefits. Acute infection encompasses the first and typically greatest peak in parasite load in human malaria infections (Miller *et al*., 1994; Eichner *et al*., 2001); subsequent waves of infected RBC abundance tend to be smaller as immune responses suppress parasite proliferation (Childs & Buckee, 2015). In experimental rodent infections, the acute phase corresponds to the greatest decline in the host RBCs (Huijben *et al*., 2010) and represents a time-frame with substantial risk of host mortality (Kamiya *et al*., 2021). Acute infection also includes large peaks in gametocyte abundance—often the largest wave of gametocyte production—and hence transmission potential in both human and rodent malaria infections (Eichner *et al*., 2001; Huijben *et al*., 2010). Data and models suggest strong selective pressures on parasite traits that govern initial infection dynamics (Huijben *et al*., 2010; Greischar *et al*., 2016, respectively). Thus, the potential costs and benefits of larger burst sizes are likely to be most pronounced in acute infection.

We investigate how host mortality and resource limitation influence the evolution of burst size and transmission investment during acute infection, using a mechanistic within-host model of rodent malaria (*P. chabaudi*) infections. Our model framework relaxes two key assumptions of previous theory (Koella & Antia, 1995): (1) that parasites grow exponentially except when removed by immune defenses; and (2) that parasite fitness is proportional to the total number of transmissible gametocytes produced. First, we omit immunity, since innate immunity can be overwhelmed by high parasite numbers, and adaptive immunity is a stronger influence on chronic rather than acute infection dynamics (Mota *et al*., 1998; Antia *et al*., 2008; Metcalf *et al*., 2011). We instead focus on resource availability, which is a major driver of acute infection dynamics and prevents parasites from sustaining exponential growth (Haydon *et al*., 2003; Mideo *et al*., 2008; Metcalf *et al*., 2011; Pollitt *et al*., 2015). Second, we allow the fitness (i.e., transmission) benefits of gametocyte production to vary with gametocyte density in line with empirical data. When gametocytes are rare, increasing gametocyte abundance can substantially increase host infectiousness, but those gains saturate when gametocytes are abundant (Paul *et al*., 2007; Huijben *et al*., 2010; Bell *et al*., 2012). We identify optimal burst size and transmission investment from simulated infections, i.e., the traits that maximize cumulative transmission potential (parasite fitness) at the end of the acute phase. Imposing an empirically-inspired minimum RBC threshold for survival, we find that the resource limitation is a stronger selection pressure on burst sizes than host mortality. We find that optimal strategies depend critically on merozoite longevity and initial RBC abundance. While our focus is on the acute phase, we find that larger burst sizes may be advantageous if parasites are able to persist within the host long after the acute phase. Our findings suggest resource limitation could be an important mechanism preventing evolution towards ever more rapid parasite proliferation.

## Materials and methods

We use a within-host model to project the impact of burst size and transmission investment on the probability of transmission to mosquitoes, summed over the lifespan of infection (cumulative transmission potential). Cumulative transmission potential has been used as a fitness metric to identify optimal traits in HIV (Fraser *et al*., 2014) and malaria infections (Greischar *et al*., 2014, 2016, 2019) and is analogous to lifetime reproductive effort in macroorganisms (Maynard Smith, 1978). To understand how resource limitation impacts cumulative transmission potential, we define the effective merozoite number within the host (*R_M_*), the average number of merozoites committed to proliferation generated by a single invading merozoite, akin to the effective reproductive number that defines the number of secondary infections produced by each infected host during an epidemic. The effective reproductive number falls below replacement level (*R_M_*= 1) as susceptible RBCs are depleted and increases past replacement level as RBCs are replenished and iRBC numbers begin to increase again. The time required for *R_M_* to increase back to one following the initial decline serves to define the end of the first acute wave of infection, though we also account for host mortality prematurely ending infection, as well as an arbitrary infection length for comparison with previous models (Greischar *et al*., 2016). Following infections over the first acute wave of infection encompasses two important aspects of the biology of pathogenic organisms. First, pathogenic organisms should be most vulnerable to extinction when susceptible individuals are depleted following an outbreak (King *et al*., 2009), and that logic should apply equally when the susceptible individuals are RBCs and the outbreak has occurred within a host. Second, the duration of infection should depend on the traits of the pathogenic organism. We then examine the sensitivity of optimal trait values to parameter choices, especially those influencing resource availability.

### Quantifying cumulative transmission potential

We assume that selection will maximize parasites’ cumulative transmission potential, infectiousness to mosquito vectors over the duration of infection. The cumulative transmission potential *f* (*ε*) is the Riemann sum of host infectiousness (*P_C_*) through the endpoint of infection (*ε*):

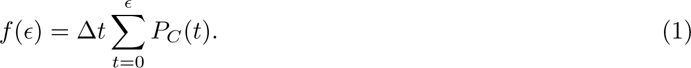

Using an empirically-derived curve from mice infected with *P. chabaudi* (Bell *et al*., 2012), we calculate the expected fraction of mosquitoes infected following a bloodmeal (*P_C_*) as a function of the abundance of mature gametocytes per microliter of blood (*G*) at each time *t* during the infection:

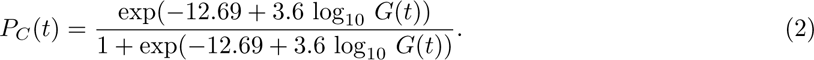

In Equation 2, when gametocytes are rare, there is a disproportionate gain in transmission success with a small increase in gametocyte abundance. Although we do not consider sex ratio explicitly, with a low number of gametocytes, the mosquito is unlikely to have both female and male gametocytes in the bloodmeal (Bell *et al*., 2012). Increasing gametocyte numbers, however, brings diminishing returns in host infectiousness.

We simulate gametocyte abundance per microliter of blood by modifying a previous mathematical model (Greischar *et al*., 2014, 2016) of within-host dynamics of the rodent malaria parasite, *P. chabaudi* (Figure 1). As before, uninfected RBC abundance (*R*) is maintained at a homeostatic equilibrium (*R^*^* = *R*(0)) in the absence of infection:

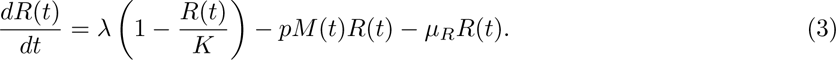

**Figure 1:**
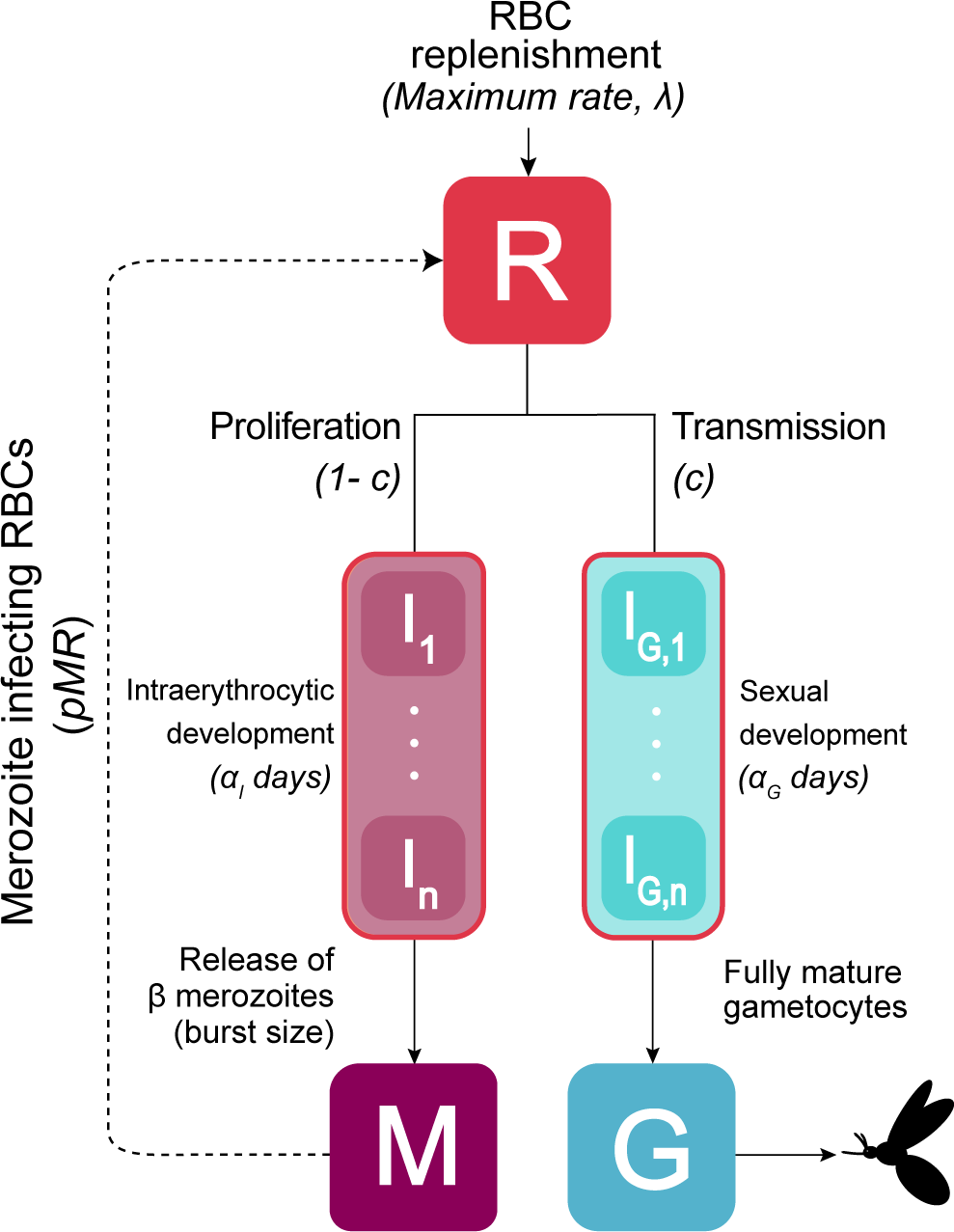
Our within-host model tracks abundance of uninfected RBCs and *P. chabaudi*-infected RBCs across different *n* stages of intraerythrocytic development. Here, RBCs (*R*) are invaded by merozoites (*M*) with a fraction 1 − *c* continuing proliferation by developing as infected RBCs (*I*) which burst to release *β* merozoites (the burst size) after a time delay of *α_I_*days. A fraction *c* (the transmission investment) of invaded RBCs commit to gametocyte development (*I_G_* class), which become fully mature gametocytes (*G*) infectious to mosquito vectors after *α_G_*days.

To maintain that equilibrium, the carrying capacity (*K*) is set to 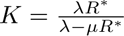 and *λ* is the maximal replenishment rate. Thus, we assume that as RBCs deviate further from the homeostatic equilibrium, there is a greater upregulation of erythropoiesis (Jakeman *et al*., 1999; Mideo *et al*., 2008). In an infection, RBCs are depleted either through intrinsic mortality *µ_R_* or by invasion of RBCs by merozoites (*M*) assuming mass action contact with an invasion rate *p*.

When RBCs are invaded by merozoites, a proportion *c*—the transmission investment—enter the *I_G_* class to develop into gametocytes (*G*). The remaining 1 − *c* of invaded RBCs continue the proliferative cycle, entering the infected RBC (*I*) class that can subsequently burst to release merozoites. Previous versions of this model assumed a fixed time delay before invaded RBCs could burst to release merozoites, and we relax that assumption by allowing the time delay to be Erlang distributed (a special form of the gamma distribution). Given a positive integer shape parameter *n*, intraerythrocytic development can be tracked with a series of *n* ordinary differential equations:

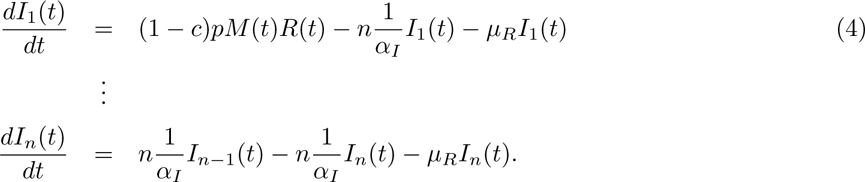

This “linear-chain trick” has individuals flowing through a series of subcompartments with the total average dwell time being *α_I_* days (Metz & Diekmann, 1991). In *P. chabaudi*, intraerythrocytic development requires one day (i.e., *α_I_* = 1 day) so that each subcompartment has a waiting time of *α_I_/n*. With only one compartment (*n* = 1), the model reduces to a standard compartmental model with exponentially distributed waiting times, while as *n → ∞* the model approaches a fixed time delay, with no variation around the mean waiting time of *α_I_*.

Infected RBCs that survive and complete intraerythrocytic development each release *β* merozoites (the burst size):

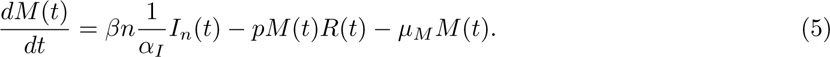

Merozoites are lost either to intrinsic mortality *µ_M_* or to successful invasion of uninfected RBCs.

To contribute to transmission success, invaded RBCs must survive the immature gametocyte stage (*I_G_*):

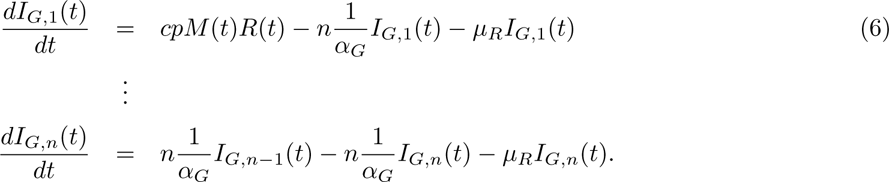

As in the *I* class, we use the linear chain trick to incorporate the gamma-distributed delay with mean of two days (*α_G_* = 2) for immature gametocytes to mature and become transmissible to the mosquito vectors (Gautret *et al*., 1996). Upon maturation, members of the *I_G_* class enter the *G* class and are assumed to be infectious to mosquitoes:

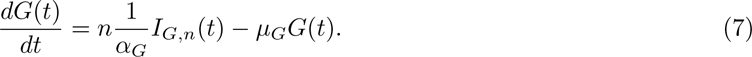

The abundance of mature gametocytes (*G*) is used to calculate the daily transmission probability (*P_C_*) and the cumulative transmission potential, *f* (eq. 1). To check whether gamma-distributed variation in developmental times alters optimal trait values, we confirm that the present model formulation gives the same optimal transmission investment (42%) found to be optimal with the analogous fixed delay model assuming an arbitrary infection length of 20 days (Greischar *et al*., 2016). To initiate the simulations, we assume a uniform age distribution of initially infected RBCs, *I*(0). All parameter values and initial conditions are listed in the Supplementary Table S1.

### Quantifying the effective merozoite number

The parasite population can only expand when each infected RBC that bursts more than replaces itself, i.e., producing more than one merozoite, committed to proliferation, that invades a susceptible RBC. To quantify how resource availability influences proliferation, we define the effective merozoite number or the *R_M_*:

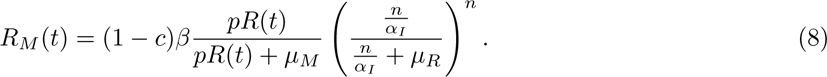

When *R_M_* is greater (less) than one, parasite numbers are expanding (declining). The (1 − *c*)*β* term is the number of merozoites emerging from infected RBCs committed to proliferation rather than gametocyte production. This value is then multiplied by the proportion of merozoites infecting a susceptible RBC rather than succumbing to intrinsic mortality, 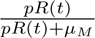, reflecting the competing rates between merozoites successfully infecting RBCs at a per-merozoite rate of *pR*(*t*) and perishing at rate *µ_M_*. Because merozoites are short-lived, with an assumed lifespan of roughly 30 minutes (McAlister, 1977), RBC depletion makes it more likely for merozoites to perish rather than infecting RBCs. We then multiply by a correction factor that accounts for the time needed for parasites to develop within the RBC. Again, there are competing rates between the parasites exiting a developmental age class (rate 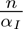) and perishing (rate *µ_R_*). Since parasites must successfully complete *n* developmental age classes comprising intraerythrocytic development, the proportion of parasites successfully maturing is raised to the *n* power. By calculating *R_M_*, the transmission probability (*P_c_*), and the cumulative transmission potential (*f*) over the infection, we can compare how resource availability influences transmission gains across a range of burst sizes and levels of transmission investment.

### Simulating the end of infection

We simulate infections by varying the burst size (*β*) from 1 to 50 and transmission investment (*c*) from 1% to 99%. Simulated infections end in one of three ways: (1) infection never establishes; (2) infection establishes but kills the host; (3) infection establishes and does not kill the host, ending either at an arbitrary time point or at the end of the acute phase (Figure 2). Infections that fail to establish are identified by calculating the *R_M_* from the initial RBC density (*R*(0)), the merozoite invasion rate (*p*), and the intrinsic merozoite mortality rate (*µ_M_*). If the parasite population cannot expand initially (*R_M_ <* 1), the infection cannot establish, but we set a higher threshold for establishment to mimic the impact of innate immunity in preventing parasite proliferation at low parasite abundance. Past work suggests that infections that expand at less than 1.5 initially struggle to increase their rates of expansion (see Fig. 1 in Metcalf *et al*. 2011), so we set the initial *R_M_*threshold to 1.5. Specifically, the inequality below must be satisfied for infections to establish:

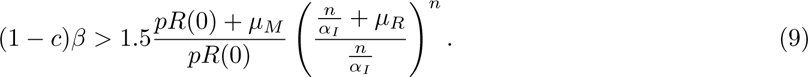

**Figure 2:**
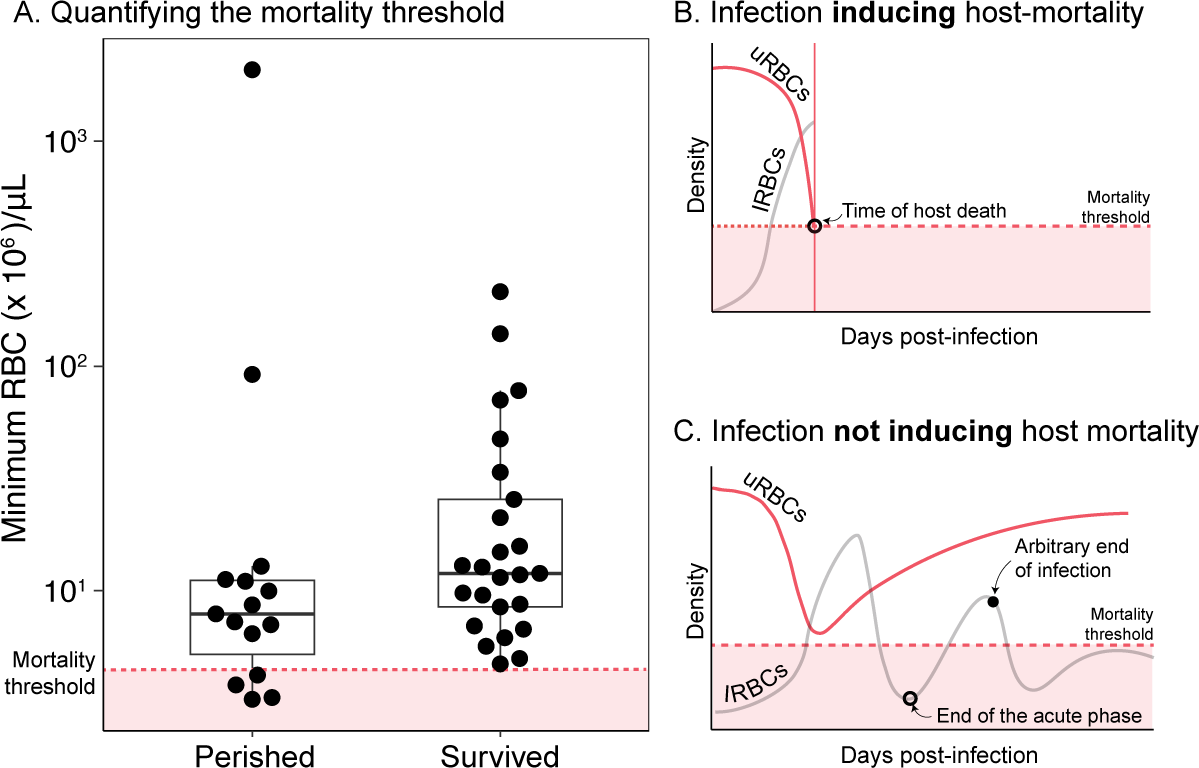
Varying the burst size (*β*) and transmission investment (*c*) can change the outcome of simulated infections. (A) Data from experimental rodent malaria infections showing the minimum uninfected RBC density for mice who either died or survived (see main text for details). The dotted red line and shaded region below indicate the minimum RBC threshold below which no mice survived, used as the mortality threshold in the within-host model. The right two panels illustrate different simulated infection endpoints with a schematic showing the abundance of uninfected RBCs (‘uRBCs’, red curves) and infected RBCs (‘iRBCs’ in light gray). (B) Of the simulated infections that establish, some lead to disease-induced mortality, where host RBCs are depleted to levels that cause catastrophic anemia and eventual death. In this case, the cumulative transmission potential (*f*) is calculated up to the time of death. (C) Other simulated infections successfully establish and do not subsequently induce host mortality. These infections are either assumed to end after the first wave of infection (open circle), or after an arbitrarily chosen amount of time has passed (closed circle), with cumulative transmission potential calculated up to the chosen endpoint (*ε*).

If the combination of burst size and transmission investment (*β* and *c*) do not satisfy Eq. 9, we set the cumulative transmission potential to 0.

Infections that establish can end either by inducing host mortality, at the end of the first wave of infected RBC abundance (acute phase), or at an arbitrarily chosen time point. The time until host mortality or the end of acute infection is determined by parasite burst size (*β*) and transmission investment (*c*), and we use an arbitrary endpoint to examine the impact the influence of infection length on optimal parasite traits. We quantify the mortality threshold based on the minimum uninfected RBC density in *P. chabaudi*-infected mice from a previous study (Mideo *et al*., 2011) (Figure 2A). In that study, mice were infected with or without treatments modifying their immune response (anti-CD4) and/or inducing RBC depletion (phenylhydrazine). Aggregating individuals based on rather they survived or perished regardless of treatment, we set the mortality threshold as the minimum RBC density of the surviving group, 6.5 × 10^5^/*µ*L. We assume host death if RBC density reaches this critical threshold and calculate cumulative transmission potential up to the time of death (Figure 2B). For infections that establish and do not induce host mortality, we first determine optimal trait combinations (burst size *β* and transmission investment *c*) based on cumulative transmission potential during acute infection. We define the length of the acute phase as the time from the beginning of the simulated infection to the time when *R_M_* = 1 following the peak infected RBC abundance (Figure 2C). When acute infection ends, we assume that proliferation ceases but we continue to calculate transmission gains from developing gametocytes as they mature, since gametocytes have been observed to persist even in hosts where proliferation has been halted (e.g., through antimalarial drugs, Bousema & Drakeley 2011).To capture any potential transmission success from gametocytes produced during the acute wave, we follow the infections to at least 100 days post-infection.

We repeat these comparisons varying the initial inoculum size, since larger (smaller) initial iRBC numbers would be expected to exaggerate (reduce) the impact of resource limitation. We also identify the optimal burst size when infections end at an arbitrary time, fixing transmission investment to the value identified as optimal for acute infection. Specifically, we calculate Equation 1 for varying infection lengths (*E* of 5, 25, and 50 days). Identifying the optimal traits for different (arbitrary) infection lengths gives context to the optima for acute infection and enables comparison with past work that assumed an arbitrary infection length (Greischar *et al*., 2016).

### Sensitivity to parameters governing resource availability

To explore how variation in parasite and host traits can influence optimal burst size, we manipulate parameters associated with *R_M_*, the merozoite mortality rate (*µ_M_*) and initial RBC abundance (*R*(0)), as well as the maximum RBC replenishment rate (*λ*). Varying each individual parameter by −/+25%, we identify changes in the optimal burst size. For comparison, 25% variation in initial RBC is consistent with experimental data from *P. chabaudi*-infected mice whose initial RBC abundance varies from 5.41 × 10^6^/*µL* to 8.57 × 10^6^/*µL* with an average of 6.85 × 10^6^/*µL* (Figure S1). When manipulating the initial RBC abundance, we keep *λ* at the original value and adjust the carrying capacity (*K*) to ensure that the homeostatic equilibrium is maintained. When manipulating *λ*, we ensure that the homeostatic equilibrium (*R^*^*) would be maintained at the same level as before (8.5 × 10^6^/*µL*) in the absence of the infection.

## Results

### Aggressive proliferation does not maximize parasite fitness during acute infection

To maximize cumulative transmission during the acute phase, the optimal strategy is for parasites to restrict their proliferation below what is required to avoid prematurely killing the host (Figure 3A). Infections fail to establish when trait combinations include either a low burst size and/or high transmission investment (light blue area, Figure 3A), i.e., when the initial effective merozoite number, *R_M_ <* 1.5. With higher burst sizes (*β*) and restrained transmission investment (*c*), infections often kill hosts by depleting uninfected RBC abundance below the lethal threshold (green lines, Figure 3A). The cumulative transmission potential at the time of host death is substantially lower than the cumulative transmission potential for trait combinations that persist longer by not inducing host mortality. Consistent with the assumptions of a transmission-virulence tradeoff, host mortality is detrimental for parasite transmission, which is terminated prematurely, typically before the parasites reach their peak parasite load.

**Figure 3:**
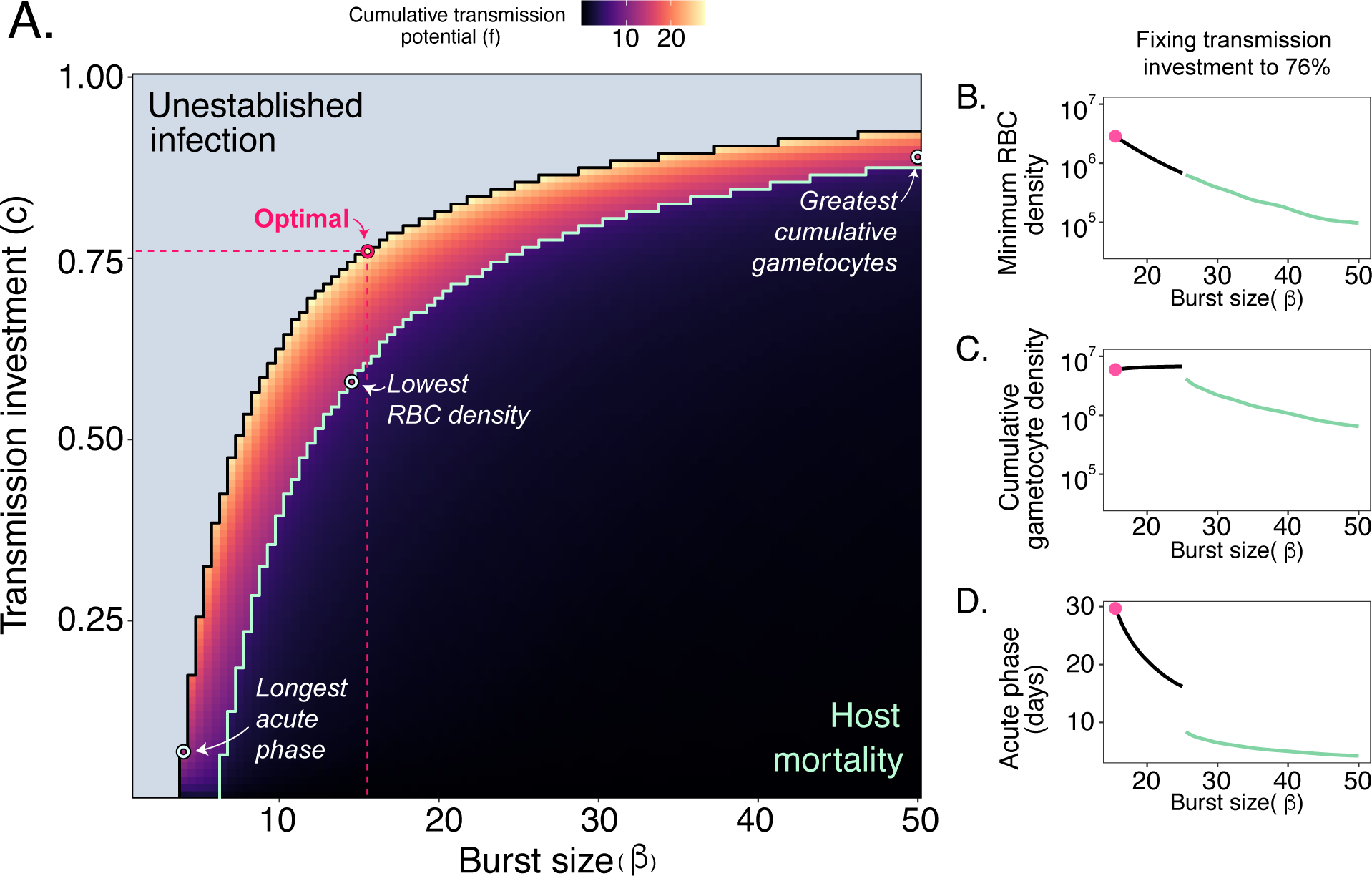
The maximum cumulative transmission potential (*f*) is achieved with an intermediate burst size that does not maximize acute infection duration, RBC depletion, or cumulative gametocyte production. (A) Different combinations of burst sizes (*β*, x-axis) and transmission investment (*c*, y-axis) yield a range of fitness outcomes (cumulative transmission potential, *f*, z-axis). The light blue area describes the parameter combinations that do not establish infections (i.e., their initial *R_M_ <* 1.5 and *f* is set to 0). The green line delineates the boundary below which *β* and *c* induce host mortality due to RBC depletion below the lethal threshold. The optimal burst size and transmission investment are *β* = 15.5 and *c* = 76%, corresponding to an acute infection phase lasting 32 days (open pink point). For comparison, open white points indicate trait combinations giving the longest acute phase, the lowest RBC density that does not induce host mortality, and the highest cumulative gametocyte production. Fixing *c* to its optimal value of 76% and varying burst size over the range that establishes infection, we show (B) the minimum RBC density achieved over the acute phase, (C) the cumulative gametocyte production over the acute phase, and (D) the duration of the acute phase. For panels B-D, the burst sizes that induce host mortality are shown in green with closed pink points to indicate the optimal burst size.

The optimal burst size and transmission investment—that is, the values that maximize the cumulative transmission potential over acute infection—are *β* = 15.5 and *c* = 76% (Figure 3A). The trait combination that induces the greatest depletion in the host RBCs occurs when *β* = 14.5 and *c* = 58%, just short of the host mortality threshold, while the combination with the greatest cumulative gametocyte production is *β* = 50 and *c* = 89% (Figure 3A). Across all parameter combinations, the longest acute duration (33.1 days) requires slow proliferation (*β* = 4, *c* = 7%), and that pattern holds across a range of initial inoculum sizes (Figure S3). Taking a cross-section of the fitness landscape by assuming a fixed transmission investment of 76%, we find that, as expected, increasing burst size leads to greater depletion of RBCs, although substantial increases in burst size are needed to induce host mortality (Figure 3B). Increasing burst size increases cumulative gametocytes produced, until the RBC depletion is so great that it abbreviates the acute phase by killing the host (Figure 3C). Host-killing trait combinations generate infections lasting less than 10 days (Figure 3D). For a given level of transmission investment (here, 76%), the optimal burst size has the longest acute phase of 29.7 days, but longer acute infections are possible with slower proliferation (lower burst sizes and transmission investment, Figure 3A). Therefore, the optimal strategy is distinct from strategies that maximize host exploitation, transmission stage production or infection length.

Critically, burst sizes could increase substantially from the optimal value without causing additional host mortality, in contrast to the assumption underpinning virulence-transmission tradeoffs, that faster proliferation generates faster host mortality. We find the identical optimal strategy when we simulate the model without a mortality threshold (Figure S2) and the same qualitative pattern—that optimal burst sizes fall below the threshold required to avoid premature host death—with different initial inoculum sizes (Figure S4). While parasite strains with higher burst sizes may produce more gametocytes during the acute phase, that need not correspond to gains in cumulative transmission potential because (1) transmission probability saturates with increasing gametocyte density (Equation 2) and (2) the acute phase is significantly shortened due to faster depletion in RBCs even in the absence of host mortality. These results show that host mortality need not be the primary mechanism restricting the evolution of faster proliferation in malaria parasites.

### Resource depletion limits optimal burst size during acute infection

To determine how resource (i.e., RBC) limitation influences optimal burst size, we simulate dynamics of suboptimal, optimal, and supraoptimal burst sizes (*β* = 14.5, 15.5, and 16.5) while assuming a transmission investment of 76%. In simulated infections, the initial effective merozoite number (*R_M_*) is the highest value, since it reflects the largest RBC abundance, prior to depletion. For all three burst sizes, RBC depletion eventually pushes *R_M_* below 1, beginning a decrease in infected RBCs abundance (dotted lines, Figure 4A). Larger burst sizes generate faster and more extreme fluctuations in effective merozoite numbers, and subsequently more extreme oscillations in gametocyte abundance and the transmission probability. After a two day lag corresponding to the time required for gametocyte development (*α_G_* = 2 days), gametocyte density and transmission probabilities (*P_c_*) also begin to decline (Fig. 4B, C). With a supraoptimal burst size (*β* = 16.5), the infection begins with a large *R_M_* (Fig. 4A), indicative of faster parasite proliferation and subsequently greater gametocyte production (Fig. 4B). While larger burst sizes generate a greater peak in gametocyte density (Figure 4B), the peak transmission probability is similar for the optimal and supraoptimal burst sizes due to the saturating relationship between transmission probability and gametocyte abundance (Figure 4C). With a burst size of 14.5, the initial *R_M_* falls below the threshold for establishment (1.5), but we include it here for comparison. Therefore, under the assumption that parasites must proliferate faster to establish in the face of innate immune defenses, it enforces a larger burst size than would otherwise be optimal. That trait combination proliferates very slowly over a very long acute phase, in contrast to the optimal and supraoptimal burst sizes. This strategy would have had a higher cumulative transmission potential than the others if it had not fallen below our threshold for establishment. Because the duration of acute infection is determined by the effective merozoite number (*R_M_*), rapid proliferation shortens the acute phase (filled points, Figure 4). Although the highest burst size shows the greatest initial increase in transmission probability, the lower, optimal burst size prolongs acute infection and hence sustains an elevated transmission probability for an extended period (Figure 4C).

**Figure 4:**
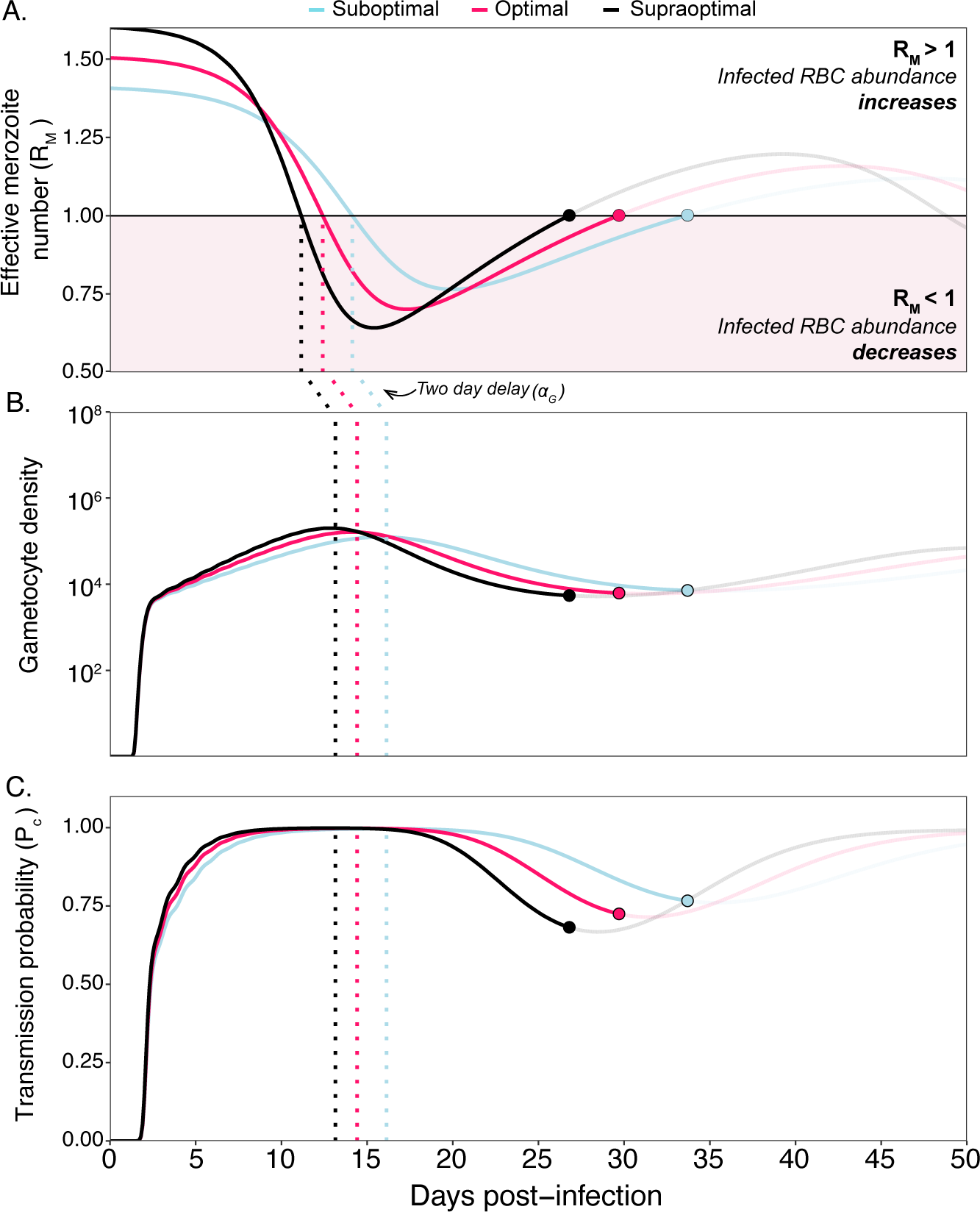
RBC depletion reduces subsequent gametocyte production and host infectiousness. The cumulative transmission potential at the end of the acute phase is determined by merozoites successfully infecting RBCs. Fixing the transmission investment (*c*) at 76% (the optimal level, Fig. 3, we simulate within-host dynamics for strains with suboptimal, optimal, and supraoptimal burst sizes (*β* = 14.5, 15.5, and 16.5, respectively). The strain with *β* = 14 does not establish (i.e., initial *R_M_ <* 1.5), but dynamics are shown for comparison. (A) When the *R_M_* —the expected number of merozoites invading and continuing the proliferative cycle—falls below 1 (pink area), the iRBCs begin their decline. (B) One developmental delay (*α_G_*= 2 days) after the iRBCs begin to decline (dotted black lines), gametocyte abundance also starts to decline. (C) That decline in gametocyte abundance reduces transmission probabilities (*P_C_*). For all panels, filled points represent the end of the acute phase for each burst size, with subsequent dynamics shown with transparent lines.

### Optimal burst size increases with the difficulty of RBC invasion

By modifying parameters that modulate ease of RBC invasion, we find broadly that larger burst sizes are optimal when RBCs are more difficult to invade, including when merozoites die faster (larger *µ_M_*, Fig. 5A) and when initial RBC abundance is lower [*R*(0), Fig. 5B]. For clarity we show optimal burst size when the transmission investment is fixed at 76%, but the same patterns hold when transmission investment is also allowed to vary (Fig. S5-S7). Increasing merozoite mortality or reducing initial RBC abundance narrows the range of trait combinations that can establish infections without inducing host mortality and also decreases the length of the acute phase (Fig. S5-S7). With a 25% higher merozoite mortality rate (i.e., shorter lifespans), we find that the optimal burst size increases to *β* = 18 from *β* = 15.5. Conversely, with a 25% lower merozoite mortality rate (i.e., longer lifespans), the optimal burst size decreases to *β* = 13.5. Increasing the initial RBC by 25% reduces the optimal burst size (*β* = 14), and when the initial RBC is decreased by 25%, the optimal burst size increases to *β* = 18.5.

**Figure 5:**
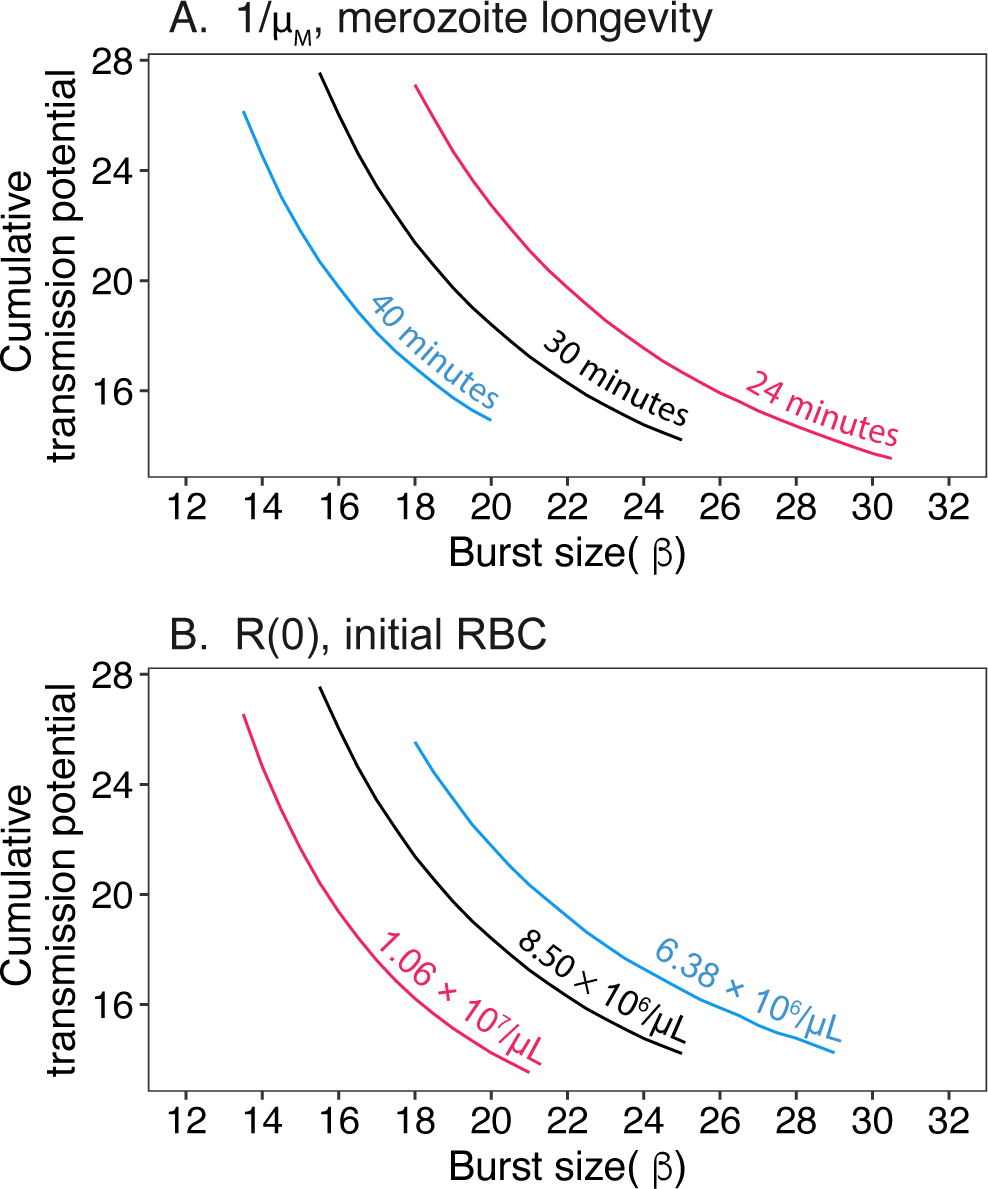
The optimal burst size decreases with ease of RBC invasion. Fixing the transmission investment to 76%, we identify the optimal burst size (*β*) while increasing (red) or decreasing (blue) each parameter by 25%. (A) The lower merozoite mortality rate (*µ_M_*, longer merozoite lifespan) selects for a lower burst size than the baseline parameter values, while the greater merozoite mortality rate (shorter lifespan) selects for larger burst sizes. With higher (lower) initial RBC abundance [*R*(0)], we find that the optimal burst is smaller (larger) than the baseline.

We also test the sensitivity of optimal burst sizes to the maximum rate of RBC replenishment (*λ*), which influences RBC dynamics but does not immediately impact the ease of merozoite invasion. Accordingly, we find that the optimal burst size remains unchanged from the original model (*β* = 15.5, Figure S8). Changes to the replenishment rate primarily alter RBC dynamics after the peak infected RBC abundance (Figure S9) and are therefore too late to influence the optimal burst size for acute infections.

### Increasing infection length reduces optimal burst size

Optimal burst size varies with total infection length, fixing the transmission investment, *c* to 76% (Figure 6). In extremely short infections (< 10 days), the best strategy is for the parasites to replicate aggressively, since infections will end before the host dies. As infection length is increased to 25 days, the optimal burst size declines well below the threshold for inducing host mortality (*β* = 25.5; red line, Figure 6A). For infections lasting 25-50 days, the optimal burst size is 15.5, the same value that maximizes cumulative transmission potential from acute infection. When infection length is intermediate (e.g., 25-50 days), either one or two peaks in transmission probability can be achieved, depending on the burst size. As in Figure 4, smaller burst sizes cause slower, gentler fluctuations in transmission probabilities over the course of infection (Figure 6). The smaller burst sizes that are optimal for infections lasting 25-50 days exhibit slower proliferation and sustain maximal transmission probabilities longer than larger burst sizes, which peak and then decline more quickly. When we allow both burst size and transmission investment to vary, we see again that the optimal burst size decreases with infection length (Figure S10). These findings suggest that intermediate burst sizes are favored in longer infections.

**Figure 6:**
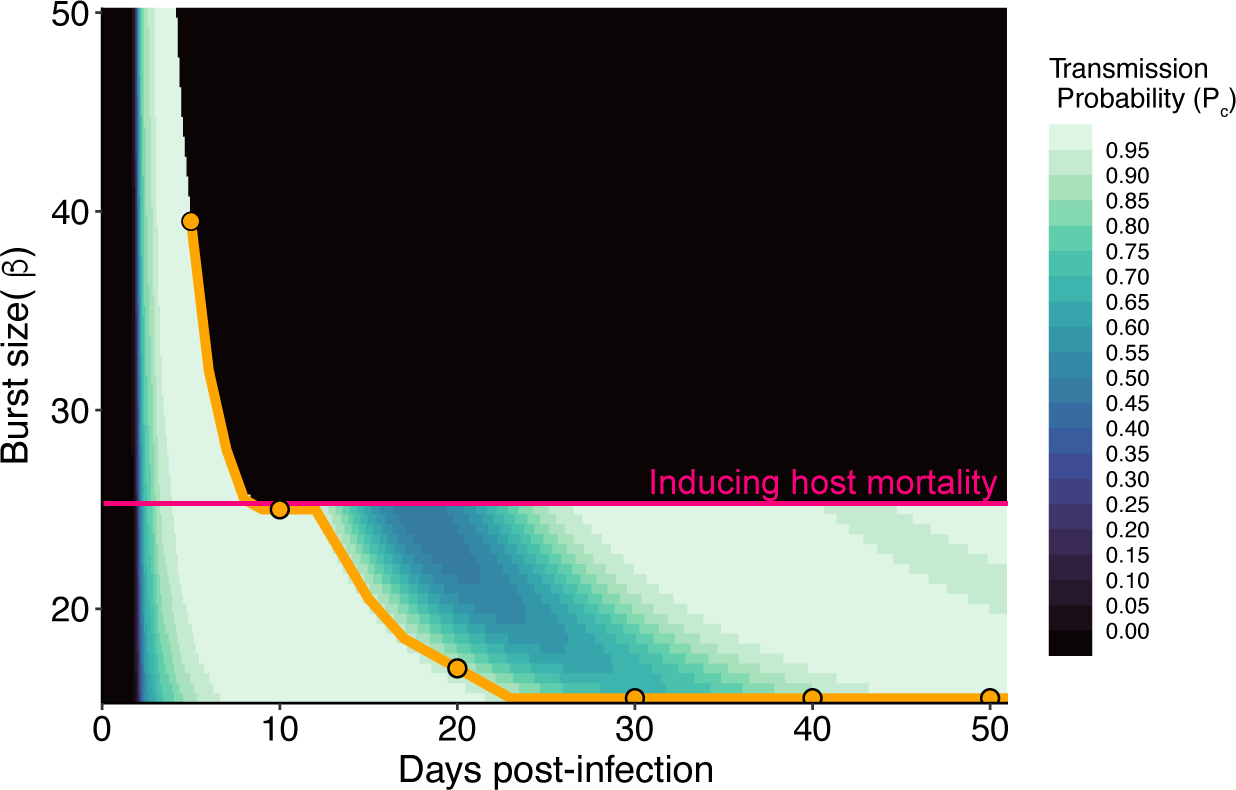
The optimal burst size (*β*) depends on the length of infection. Varying burst size (y-axis) leads to differences in the dynamics of transmission probability (*P_C_*, z-axis) over the entire infection (x-axis). Optimal burst sizes (orange line and circles) for a given infection length (indicated by position on x-axis) decrease with total infection length before increasing to a value just below what would induce host mortality (red line). Burst sizes that fall above the dashed red line deplete RBCs to the lethal threshold but can still be favored in very short infections (< 10 days), which end before RBCs reach that threshold. Transmission investment (*c*) was set to 76%.

## Discussion

A tradeoff between virulence and transmission is often assumed to prevent parasite evolution towards ever greater rates proliferation (Anderson & May, 1982; Frank, 1996; Alizon *et al*., 2009). Yet low case fatality rates in many pathogenic organisms raise the question of what forces besides disease-induced mortality could influence the evolution of proliferative capacity (reviewed in Bull & Lauring, 2014). One possible constraint on the evolution of faster proliferation is resource limitation, which is both ubiquitous and appears to play a critical role in shaping the dynamics of acute infection (Haydon *et al*., 2003; Metcalf *et al*., 2011, 2012). Nonetheless, within-host models often assume that pathogenic organisms grow exponentially and are only limited by immune clearance (Koella & Antia, 1995; King *et al*., 2009). We use a model of malaria infections— which meet the key assumptions for a virulence-transmission tradeoff (Mackinnon & Read, 2004)—to show that resource (RBC) availability restricts parasite proliferation to a far greater degree than infection-induced host mortality. Both the burst size and transmission investment influence the rate at which RBCs are depleted which directs the ‘tempo’ of the acute phase, notably the timing and duration of acute infection. The dynamics of parasite proliferation during the acute phase is predicted to have a disproportionately large impact on parasites’ fitness (Greischar *et al*., 2016), and we find that maximizing transmission during this initial phase of the infection necessitates an intermediate burst size. Our findings challenge the conventional wisdom that transmission-virulence tradeoffs are the major mechanism limiting the evolution of faster within-host proliferation.

Previous theory suggested that parasites could maximize transmission by increasing intrinsic growth rates (akin to burst size in the present study) while also increasing transmission investment to avoid reaching lethal densities within the host (Koella & Antia, 1995). That model assumes that (1) parasites proliferate exponentially, without resource limitation, and that (2) selection will maximize cumulative gametocyte production. Our model instead predicts that intermediate burst sizes maximize transmission by incorporating key details of within-host ecology likely to be common across pathogenic organisms. First, we allow for resource limitation (depletion of susceptible RBCs) as a result of proliferation. Second, we assume selection will maximize cumulative transmission potential, rather than the production of transmissible gametocytes. There is an important distinction between these two fitness metrics because the probability of transmission must necessarily saturate with increasing gametocyte numbers. The assumption of diminishing returns in transmission probability was inspired by empirical patterns in malaria infections (Paul *et al*., 2007; Huijben *et al*., 2010; Bell *et al*., 2012) but is also reflected in general models linking within-host dynamics to epidemic spread (e.g., King *et al*., 2009). We find that the burst size and level of transmission investment that maximize cumulative gametocyte production over the acute phase are different from the combination maximizing cumulative transmission potential (Figure 3A). Due to RBC availability limiting parasite expansion, our model imposes a significant cost on fast proliferating trait combinations as increases in gametocyte production do not translate to proportional transmission gains. Fast proliferation also greatly shortens acute infection, hastening the time at which parasite populations ebb and, presumably, become most vulnerable to immune clearance or to stochastic extinction (King *et al*., 2009). As a result of incorporating resource limitation and saturating transmission gains, our model readily recovers the pattern that intermediate, rather than maximal, proliferation rates are optimal.

Malaria infections seem to exhibit far more restrained transmission investment than initially expected (Taylor & Read, 1997). Our work supports the idea that extremely high levels of transmission investment are maladaptive as parasites will struggle to establish an infection, in line with past modeling studies showing that allocating resources for onward transmission, particularly early in infection, can slow proliferation and delay gametocyte production, hindering future transmission success (Koella & Antia, 1995; Mideo *et al*., 2008; Greischar *et al*., 2016, 2019). Nonetheless, the optimal level of transmission investment we identify here (76%) is substantially higher than that predicted for *P. chabaudi* by previous studies (e.g., 42%, Greischar *et al*., 2016). That study assumed a burst size of 10 and showed that larger burst sizes enable greater transmission investment. Since we recover an optimal burst size of 15.5 (Fig. 3) that enables greater transmission investment than previously predicted. That optimal burst size is somewhat higher than the range that has been reported for *P. chabaudi* (2-14, Mideo *et al*., 2011), and we find that other aspects of ecology, especially low initial inoculum sizes, can lead to substantially higher optimal burst sizes (> 40) (Fig. S4).One possible explanation for why we predict larger than observed burst sizes may rest with a key assumption of our model, that RBCs are the limiting resource for parasites. Other nutrients can be important limiting resources for parasites (Wale *et al*., 2017), and incorporating the dynamics of multiple resources may be important for understanding the bounds on burst size evolution. Mapping the constraints on burst sizes is necessary to make robust predictions about transmission investment.

Over-investing in transmission is predicted to be even more costly when parasites face host immunity and competitive displacement by other strains (McKenzie & Bossert, 1998; Mideo & Day, 2008; Greischar *et al*., 2016, 2019). The impact of infection length provides a distinct, indirect route by which immunity and coinfections could impact the evolution of proliferation rates, adding to past theory showing that immune defenses and competition from a coinfecting strain can provide strong, direct selection for aggressive proliferation (Mideo & Day, 2008; Klein *et al*., 2014; Greischar *et al*., 2016, 2019). Adaptive immunity plays a crucial role in determining the rate at which malaria parasites are fully cleared (Klein *et al*., 2014; Childs & Buckee, 2015). For example, in *P. chabaudi* infections, previously-infected mice experience shorter infections compared to immunologically naive hosts (Jarra & Brown, 1985; Mackinnon & Read, 2003). Additionally, coinfections with different malaria strains and species are common (Bruce *et al*., 2000; Bruce-Chwatt *et al*., 1962; Daubersies *et al*., 1996; Farnert *et al*., 1997) and likely to affect the total infection length (Childs & Buckee, 2015). As a result, malaria infections tend to be shorter in regions where malaria is endemic, meaning hosts are repeatedly exposed to malaria infections, and where strains turnover rapidly due to displacement by coinfecting strains (Bruce-Chwatt *et al*., 1962; Snounou *et al*., 1997; Galatas *et al*., 2018). If the total infection length is relatively short (but not so short that infections end before hosts die), we find that the optimal strategy involves restricting burst size. The strategy holds regardless of how we calculate the cumulative transmission potential: either 25-50 days post-infection (Figure 6A) or using the *R_M_* to quantify the length of the acute phase (Figure 3A). Our model suggests that in high transmission environments, parasites will be under strong selection to maximize transmission during the initial, acute phase of the infection, resulting in indirect selection on transmission investment and burst sizes that could run counter to direct selection for aggressive proliferation.

From our sensitivity analysis, we find that the optimal burst size for acute infection increases as we reduce the ease with which merozoites contact and invade RBCs. Specifically, the merozoite mortality rate (*µ_M_*) constrains the odds that merozoites will invade RBCs before perishing. We assume a baseline average longevity of 30 minutes, near the upper limit the inferred merozoite lifespan for another murine parasite, *P. berghei* (McAlister, 1977). We find that even a 25% increase in mortality rate, corresponding to only a 6-minute decrease in average longevity, increased the optimal burst size to 17 from 14.5 (Figure 5A). Examining merozoite longevity directly is challenging, but experimental work to date suggests merozoite lifespans are short in *Plasmodium spp.*, on the order of minutes (McAlister, 1977; Boyle *et al*., 2010). Uncertainty in empirical estimates of merozoite longevity could represent an important limitation on predictions for optimal burst size. That uncertainty could also mask variation across and within *Plasmodium spp.* that may represent an important driver of variation in burst sizes. Beyond differences in intrinsic mortality of merozoites, immune effectors can target merozoites (Costa *et al*., 2011), imposing greater mortality rates and potentially selection for larger burst sizes. The impact of innate and adaptive immunity fluctuates over the course of the infection (Kamiya *et al*., 2020), so that viability of merozoites may change drastically over the course of infection. Burst sizes are thought to be variable over the course of infection (Mideo & Reece, 2012), and our results raise the possibility that such variation could be an adaptive response to temporal changes in merozoite longevity.

Of the environmental variables that have been studied *in vitro*, temperature is known to affect merozoite lifespans. At 37*^◦^*C, the half-life of *P. falciparum* merozoite was observed to be only 8 minutes, while merozoite half-life nearly doubles to 15 minutes at room temperature (20*^◦^*C) (Boyle *et al*., 2010; Kumar *et al*., 2017). In malaria infections, febrile conditions can cause the host’s body temperature to reach 41*^◦^*C, suggesting even shorter merozoites lifespans and concomitant reductions in proliferative capacity (reviewed in Boyle *et al*., 2013). If we assume merozoite mortality varies throughout the infection, the optimal burst size may be higher or the optimal transmission investment lower than our predicted optimal values as parasites hedge their bets against a hostile environment. Our findings also suggest that host temperature may be a possible driver of the immense diversity we see in the burst size of the *Plasmodium* genus. Thermoregulatory mechanisms differ significantly between mammalian, reptilian, and avian hosts, presenting unique environments—and selective pressures—for different malaria species (Prinzinger *et al*., 1991; Seebacher & Franklin, 2005; Clarke & Rothery, 2008). Certain hosts harbor environments with more extreme temperature fluctuations, including ectothermic hosts that are unable to metabolically regulate their body temperatures (Seebacher & Franklin, 2005) and bats, which can exhibit significant variation in body temperature due to their ability to fly and enter torpor (Fumagalli *et al*., 2021). We predict that larger burst sizes represent a possible mechanism for parasites to adapt to transiently high temperatures, ensuring that some merozoites infect RBCs despite greater merozoite mortality. Adaptation to temperature extremes could be one mechanism to explain for the extraordinary burst size in certain malaria species that infect reptilian hosts, such as *P. giganteum*, where burst sizes can exceed 100 (Schall, 1990).

Similarly, changes to initial RBC availability should influence optimal burst sizes including the RBC range across hosts. Interestingly, with low initial RBCs, the optimal strategy shifts to greater burst sizes (Figure 5B) going against the intuition that the parasites should restrict proliferation to prevent over-exploitation. Because the effective merozoite number (*R_M_*, Eq. 8) depends on abundance of uninfected RBCs, reduced RBC density should select for parasites that produce more merozoites to increase the likelihood of infecting a rare resource. As with host body temperature, initial RBC availability represents a potential driver of burst size variation across the *Plasmodium* genus. Even within a host species prior to infection, there is considerable variation in the RBC availability (e.g., Figure S1). To explore the interspecific variation in burst size, allometric relationships can inform comparisons between host traits and parasite burst size, including between host body mass and average RBC density. For example, within mammal taxonomic groups, there is a negative relationship between RBC count and body mass, with larger mammals having fewer, but larger RBCs (Promislow, 1991). We predict that larger mammals with lower RBC availability should be exploited by *Plasmodium* species with higher burst sizes. However, body size also has confounding relationships with thermoregulation, necessitating careful consideration in disentangling the effect of body mass on burst size. Malaria species that exhibit different burst sizes, even when exploiting the same host (e.g., *P. chabaudi* and *P. berghei*), could be helpful for disentangling the impact of host versus parasite traits on the evolution of burst size.

Finally, the rate at which malaria parasites proliferate should affect the total infection length. Previous empirical and theoretical work has shown that greater within-host proliferation is associated with lower host recovery rates, thus prolonging the total infection length (Mackinnon & Read, 2003, 2004; Klein *et al*., 2014). Our analysis suggests that a high burst size does not maximize transmission during the acute infection but— in the context of host immunity—could greatly increase persistence of the parasite. As total infection length increases, we find that the optimal burst size decreases which suggests there is a tradeoff between maximizing the transmission potential over acute infection versus the chronic phase. This tradeoff raises a crucial question: does the optimal burst size change throughout the season with certain conditions favoring early transmission or prolonged persistence? The genetic diversity in human malaria parasites (*P. falciparum*) fluctuates through the season (Zwetyenga *et al*., 1999; Adjah *et al*., 2018), perhaps indicative of fluctuating selection for distinct life-histories at different points in the transmission season.

Distinct from the impact of infection length, the opportunities for transmission will vary seasonally. Since our model focuses on within-host dynamics, we assume that mosquito vectors are always equally available, but mosquito populations fluctuates in response to seasonal temperature and rainfall (Eikenberry & Gumel, 2018). During dry seasons, when transmission is rare, we would expect long-term persistence to carry a greater selective benefit than maximizing transmission at a shorter time-scale. We also implicitly assume that the blood-stage infection only occurs in immunologically naive hosts, and that there are no limitations on the availability of new susceptible hosts. However, theory predicts epidemiology imposes important selection on within-host traits (Day, 2003), like burst size and transmission investment (the latter examined in Greischar *et al*., 2019). For example, in an expanding epidemic, the ‘age’ of infection skews younger such that traits maximizing transmission during early, acute phase are favored. In that case, our model suggests an intermediate burst size may be favored. In addition to resource limitation, burst size may be restricted in finite host populations given past theory showing that faster-proliferating strains can rapidly burn through susceptibles, greatly increasing the likelihood of extinction (King *et al*., 2009). Thus, while we find that resource limitation is a potent source of selection against aggressive proliferation, ecology outside the host is likely to play an important role.

The optimal proliferation strategy is further complicated by the possibility that burst size may be adaptively plastic (flexible based on the environment) with parasites adjusting burst size in response to changes in the within-host environment (Mideo *et al*., 2011; Birget *et al*., 2019). Our results suggest that resource availability represents a fundamental aspect of the within-host environment that parasites would be expected to sense and adjust their proliferation accordingly. In support of that idea, recent work suggests that two human malaria species use nutrient availability to modulate their burst sizes *in vitro* (Stürmer *et al*., 2023), and it has been known for decades that malaria parasites adjust their transmission investment in response to the within-host environment (reviewed in Carter *et al*., 2013). Parasites may also be able to sense the environment outside the host, though the mechanisms remain unclear, since malaria-infected canaries become more infectious after being fed upon by uninfected mosquitoes (Cornet *et al*., 2014). Taken together, there is potential for parasites to employ adaptively plastic strategies, but much depends on whether parasites can sense environmental change and at what ecological scale.

By highlighting the mechanisms beyond host mortality, our study provides potential avenues for understanding the immense diversity in the life histories of the *Plasmodium* genus and the various strategies employed to exploit host resources. Beyond malaria parasites, our models could be adapted to other disease systems especially where there is a trade-off between maximizing transmission during acute infection or prolonged persistence in the chronic phase. By explicitly linking the transmission probability to the RBC availability, our model allows us to vary the parasite traits that govern proliferation and identify how different strategies impact parasite fitness. Our work demonstrates that incorporating detailed processes of within-host ecology—especially the ubiquitous constraint of resource limitation—can advance understanding of the checks on parasite evolution.

## Author contributions

All authors contributed in developing both the theory and methods in the paper. All authors contributed to the writing of the final manuscript. DP analyzed the data and performed the computations.

## Acknowledgements

The authors would like to thank Lauren Childs, Nicole Mideo, Aidan O’Donnell, Petra Schneider, Sarah Reece, Kayla Zhang, Weixin Du, and Martina Morelli for helpful discussion.

The authors declare no conflict of interest.

## Data Availability Statement

The data inspiring the host mortality threshold are in the process of being added to a publicly available server and are currently provided as a supplemental file. All the code for the project are available on https://github.com/greischarlab/burstsize

## Supplementary Material

**Table S1:** Parameter values, units, and sources.

| Parameter | Value | Source |
| --- | --- | --- |
| $R^*$ (red blood cell count at homeostasis) | $8.5 \times 10^6$ cells/μL | Savill *et al.* 2009 |
| $\lambda$ (maximum new red blood cells) | $3.7 \times 10^5$ RBCs/μL/day | Savill *et al.* 2009 |
| $\mu_R$ (red blood cell mortality rate) | 0.025/day | Miller *et al.* 2010 |
| $p$ (max. per merozoite invasion rate) | $4 \times 10^{-6}$/day | Mideo *et al.* 2008* |
| $\alpha$ (blood-stage delay) | 1 day | Landau & Boulard 1978 |
| $\alpha_G$ (gametocyte delay) | 2 days | Gautret *et al.* 1996 |
| $n$ (shape parameter) | 100 | Assumed |
| $\mu_M$ (merozoite mortality rate) | 48/day | used in Hetzel & Anderson 1996,<br>McAlister 1977 |
| $\mu_G$ (gametocyte mortality rate) | 4/day | Gautret *et al.* 1996† |
| $I(0)$ (initial infected RBCs) | 4385.965 cells/μL | modified from Greischar *et al.* 2016 |
| $I(0)$ (low inoculum) | 43.85965 cells/μL | Greischar *et al.* 2016 |
| $I(0)$ (high inoculum) | 438,596.49123 cells/μL | modified from Greischar *et al.* 2016 |
|  |  | * within realistic range |
|  |  | † length of most infectious stage |

**Figure S1:**
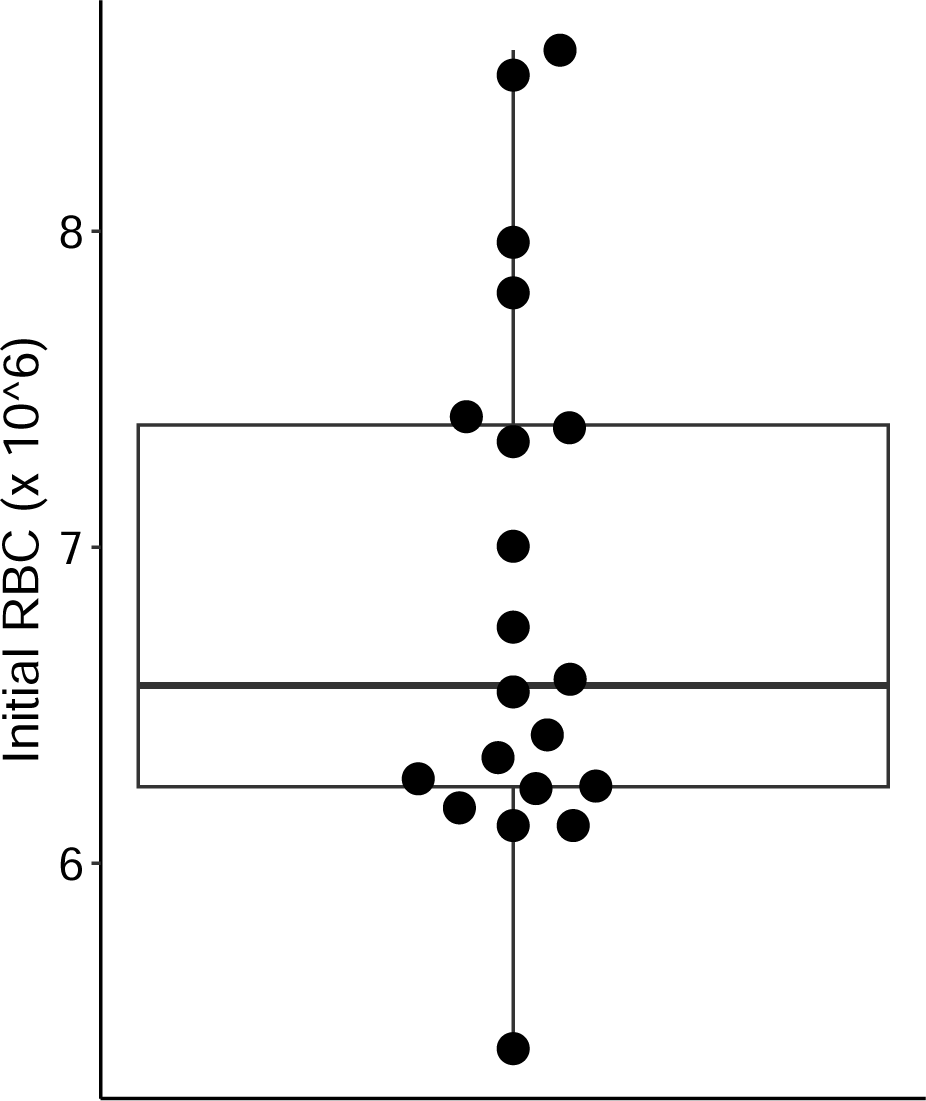
The initial RBC density of the laboratory mice 5 days before being infected with *Plasmodium chabaudi*. We exclude treatment groups that were given phenylhydrazine. The average initial RBC density ranges from 5.41 × 10^6^ × 10^6^/*µL* to 8.57 × 10^6^/*µL*, with an average of 6.85 × 10^6^/*µL*.

**Figure S2:**
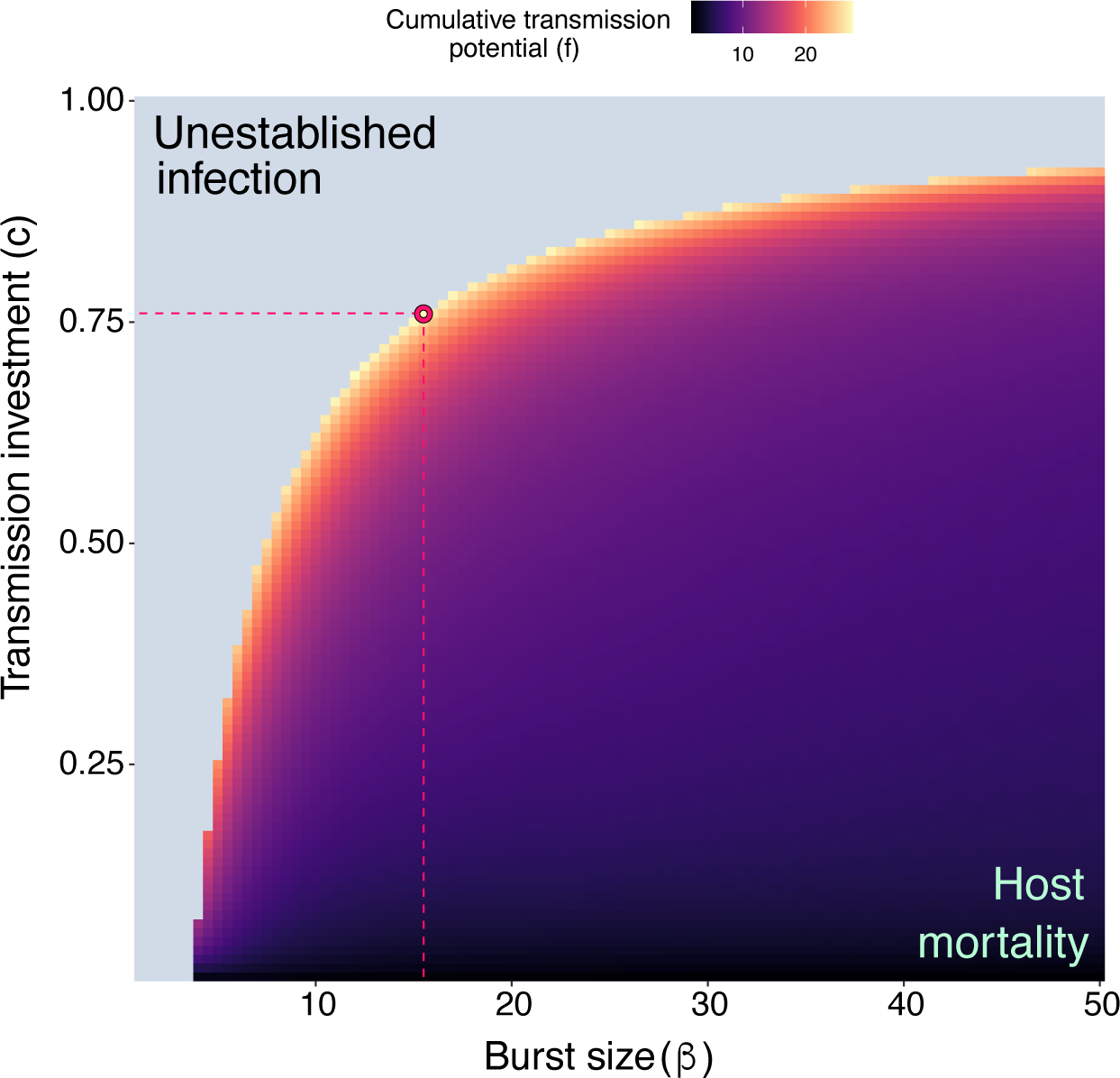
Removing the mortality threshold, we find the optimal burst size (*β*) and transmission investment (*c*) that maximize the cumulative transmission potential are the same as if there were host mortality (Figure 3A) The area in blue on the left side describes the parameter combinations that do not establish infections (*R_M_* (0) < 1.5). The pink point represents the optimal burst size of 15.5 and transmission investment of 76%.

**Figure S3:**
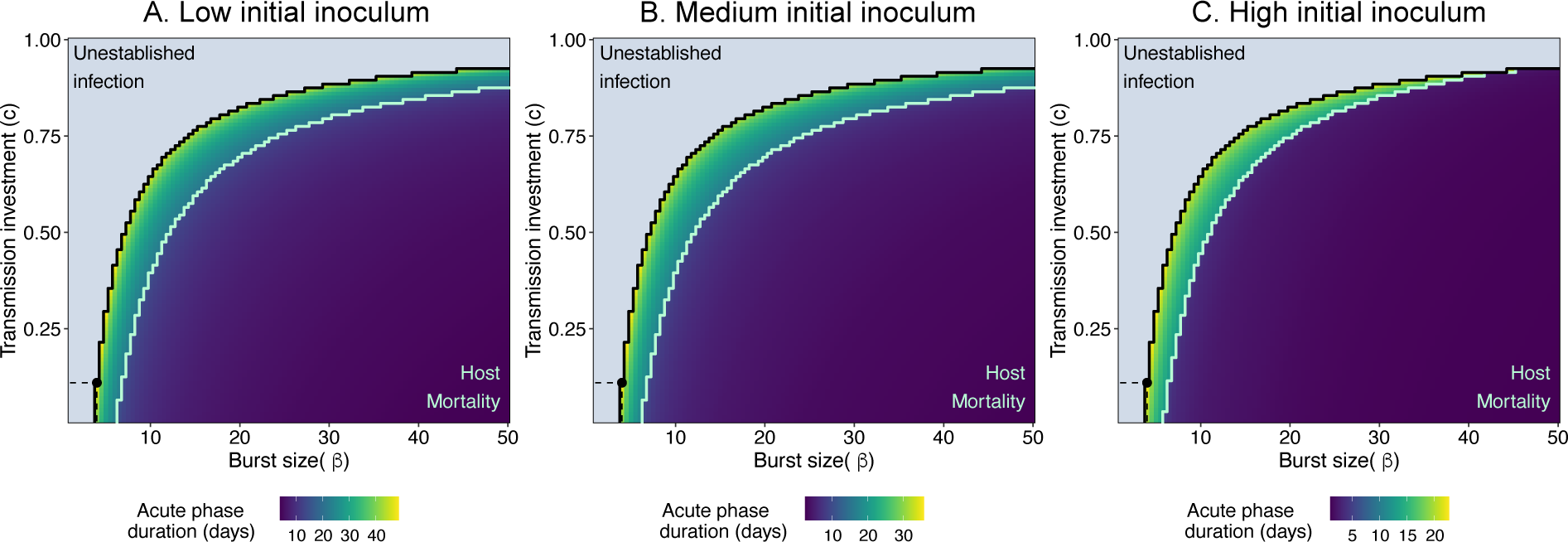
The duration of the acute phase (in days) varies with the burst size (*β*) and transmission investment (*c*) across different initial inocula: (A) 43.85965 parasites/*µL*, (B) 4,385.96491 parasites/*µL*, (C) 438,596.49123 parasites/*µL*. We find that the acute phase is longest at parameter combinations that are at the edge of not establishing infection. As burst size increases, acute infection duration is shorter. The area in blue at the left and top indicates parameter combinations that do not establish infections (*R_M_* (0) < 1.5) and the cumulative transmission potential (*f*) is set to 0. The green line indicates the boundary where the *β* and *c* induce host mortality due to catastrophic loss of host RBCs. The strain with the longest acute infection duration has a *β* = 4 and *c* = 7% across all inoculum sizes with the acute phase lasting (A) 44.5 days, (B) 33.1, and (C) 21.3 days (black point).

**Figure S4:**
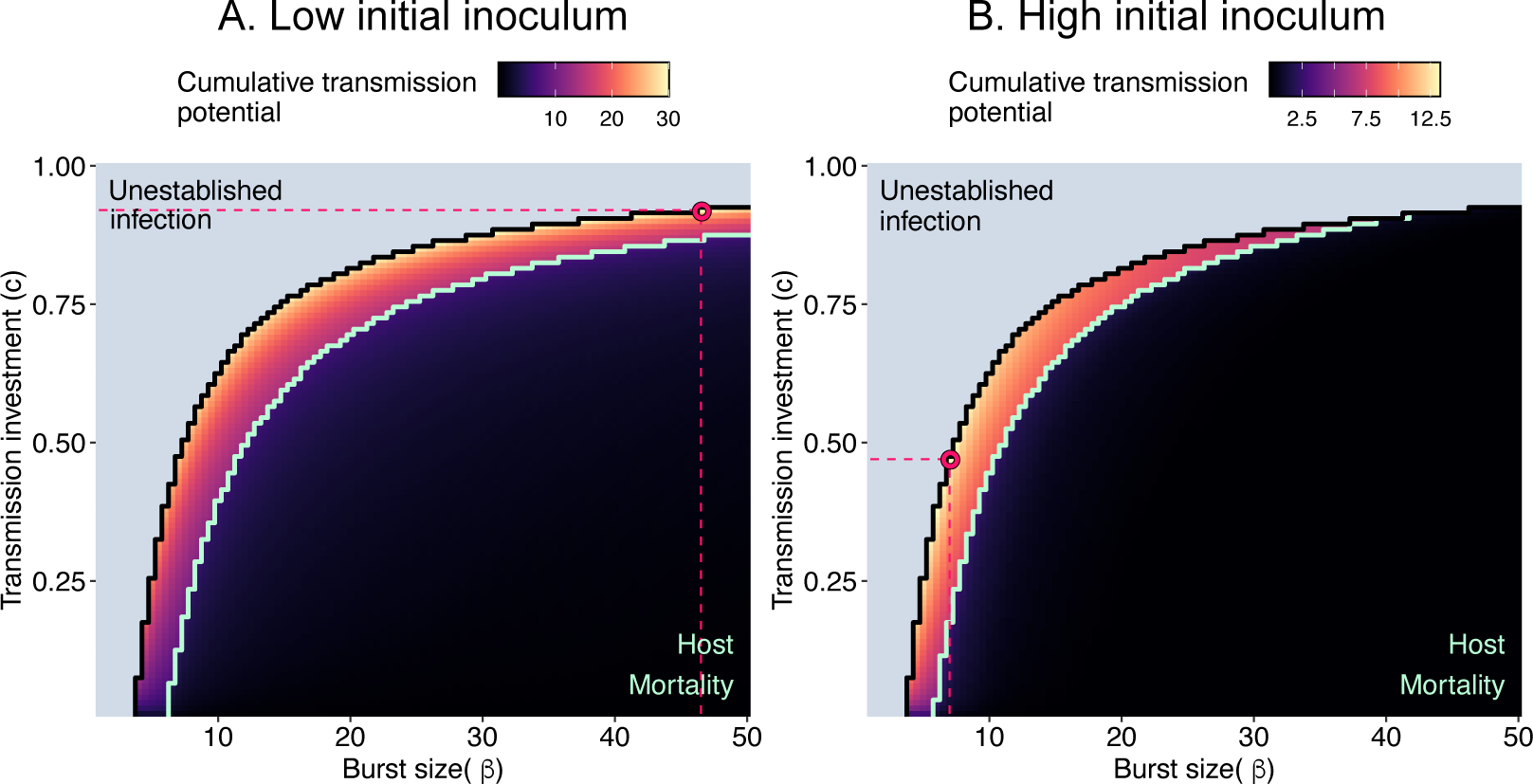
The optimal strategy maximizing cumulative transmission potential (*f*) at the end of the acute phase changes with different initial numbers of infected RBCs (inoculum size). (A) The low initial inoculum (43.85965 parasites/*µL*) has an optimal burst (*β*) of 46.5 and an optimal transmission investment (*c*) of 92% (pink point) and (B) the high initial inoculum (438,596.49123 parasites/*µL*) has an optimal *β* = 7 and *c* = 47% (pink point). Regardless of the initial inoculum, the optimal strategy is not the maximum burst size that does not induce host mortality. The area in blue on the left and top indicates parameter combinations that do not establish infections (*R_M_* (0) < 1.5), where cumulative transmission potential (*f*) is set to 0. The green line indicates the boundary where the *β* and *c* induce host mortality due to catastrophic loss of host RBCs.

**Figure S5:**
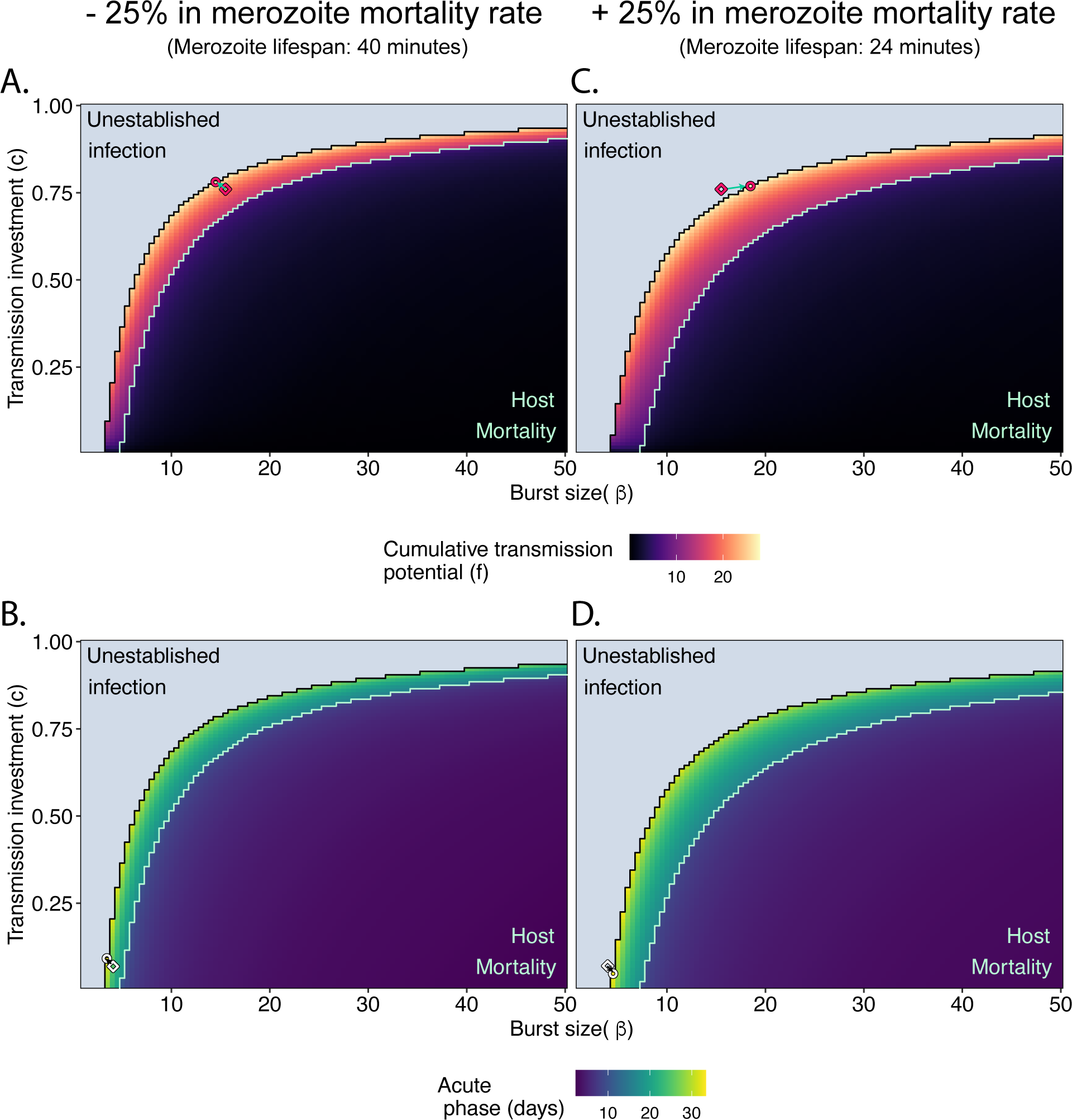
The optimal burst (*β*) and transmission investment (*c*) change when the merozoite mortality rate (*µ_M_*) is changed by +/− 25%. (A) When merozoite mortality is decreased by 25%, increasing merozoite longevity, the optimal strategy (pink circle) is *β* = 14.5 and *c* = 78%. (B) The strain with the longest acute infection duration (white point) is when *β* = 3.5 and *c* = 9% with an acute phase of 32.3 days. (C) When merozoite mortality is increased by 25%, decreasing its longevity, the optimal strategy is *β* = 18.5 and *c* = 77%. (D) The longest acute phase is when *β* = 4.5 and *c* = 5% with an acute phase of 33.4 days. The area in blue at the left and top indicates parameter combinations that do not establish infections (*R_M_* (0) < 1.5) and the cumulative transmission potential (*f*) is set to 0. The green line indicates the boundary where the *β* and *c* induce host mortality due to catastrophic loss of host RBCs. The diamonds represent the optimal strategy from the original model (*β* = 15.5, *c* = 76%) and the arrows point to the the circles that represent the new optimal *β* and *c*.

**Figure S6:**
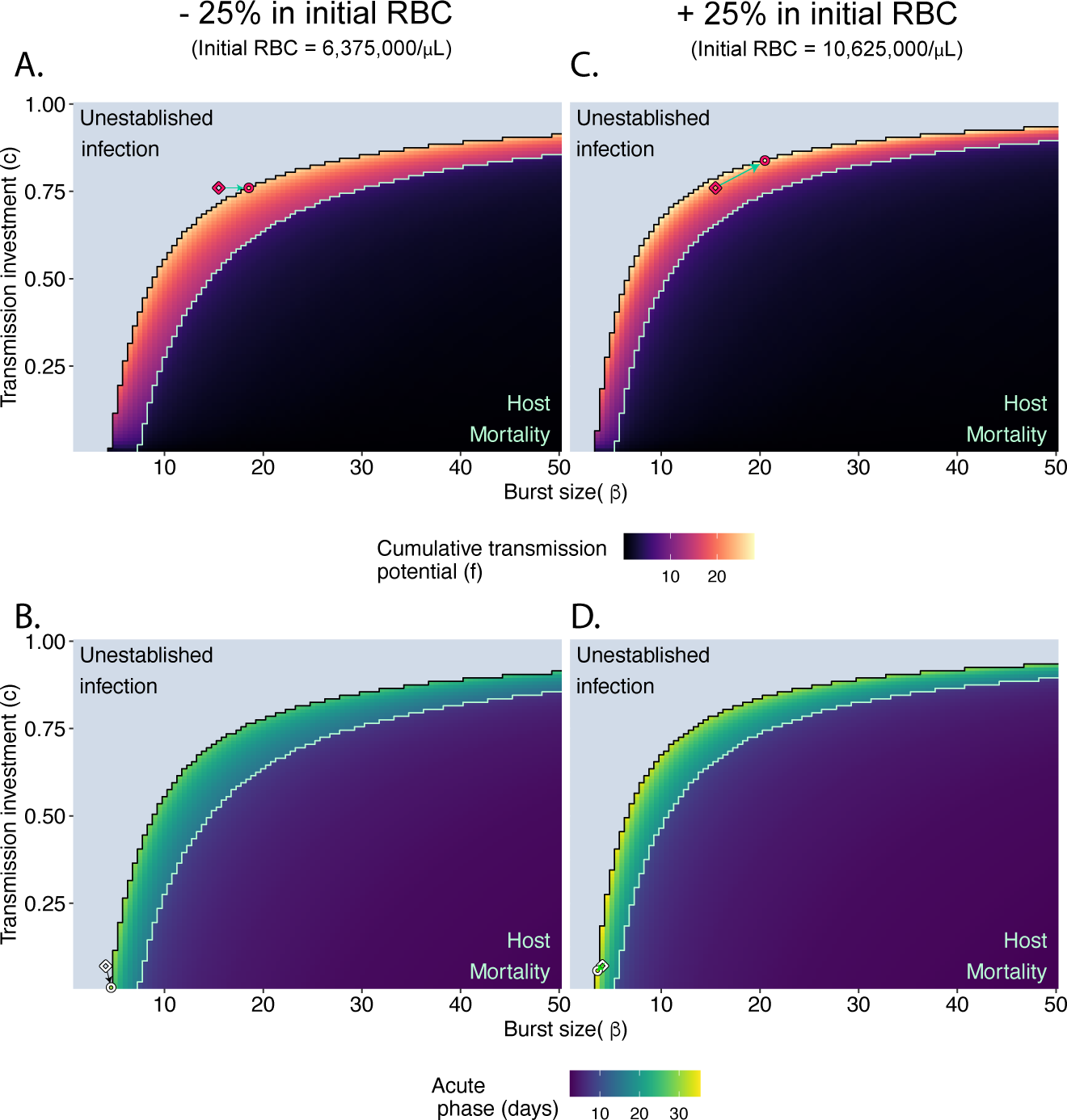
The optimum burst (*β*) and transmission investment (*c*) shift when the initial RBCs (*R*(0)) is changed by +/− 25%. (A) When initial RBCs is decreased by 25%, the optimal strategy (pink circle) is *β* = 18.5 and *c* = 76%. (B) The strain with the longest acute phase (white point) is when *β* = 4.5 and *c* = 1% with an acute phase of 30.5 days. (C) When the initial RBCs is increased by 25%, the optimal strategy is *β* = 20.5 and *c* = 84%. (D) The longest acute phase is when *β* = 3.5 and *c* = 6% with an acute phase of 35.4 days. Colors and points as in Fig. S5.

**Figure S7:**
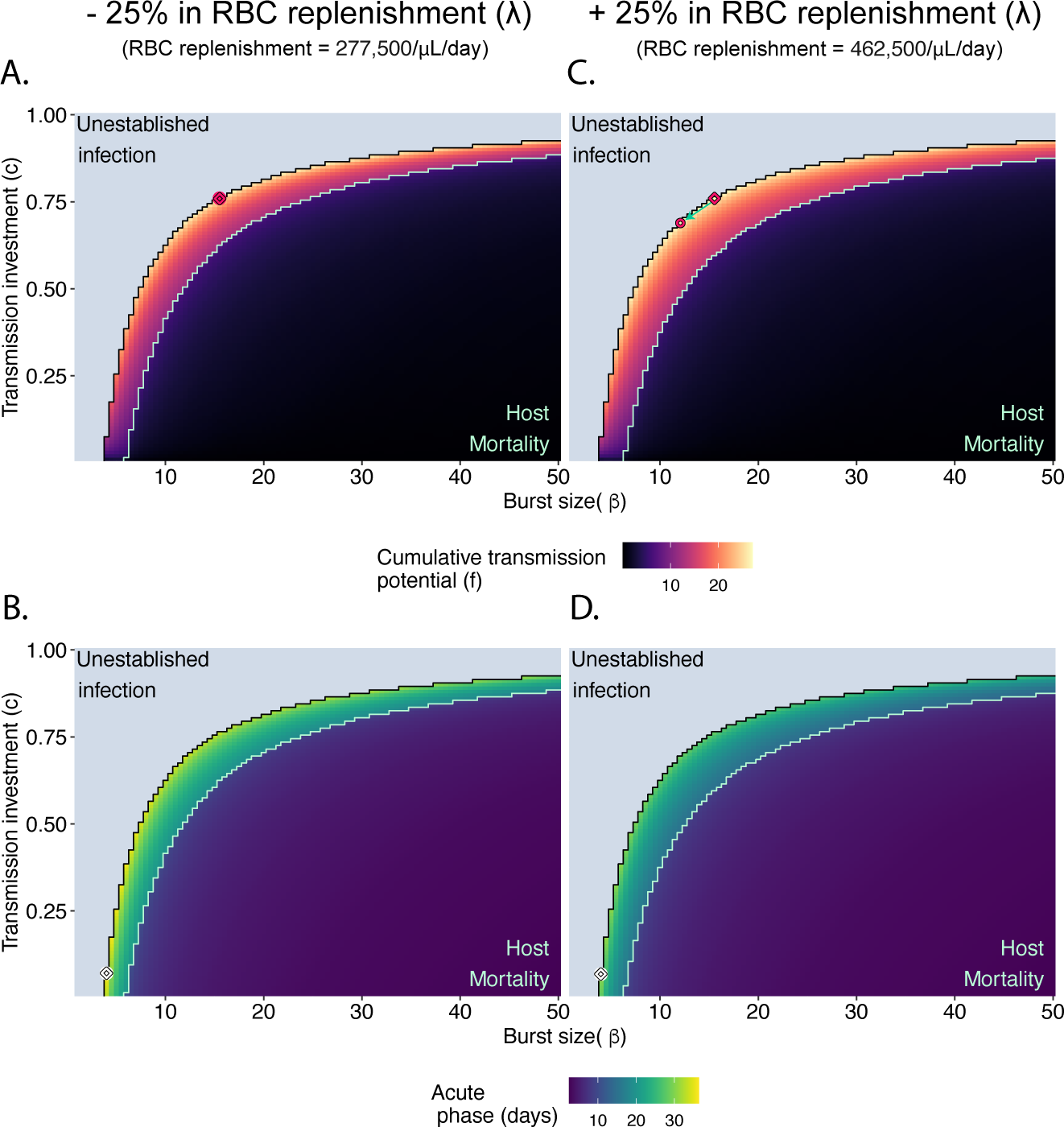
The optimum burst (*β*) and transmission investment (*c*) is nearly identical when the maximum RBC replenishment rate (*λ*) is changed by +/− 25%. (A) When *λ* is decreased by 25%, the optimal strategy (pink circle) remains the same (*β* = 15.5 and *c* = 76%). (B) The strain with the longest acute phase (white point) is when *β* = 4 and *c* = 7% with an acute phase of 36.4 days. (C) When *λ* is increased by 25%, the optimal strategy shifts slightly to *β* = 12 and *c* = 69%. (D) The longest acute phase is when *β* = 4 and *c* = 7% with an acute phase of 31.2 days. Colors and points as in Fig. S5.

**Figure S8:**
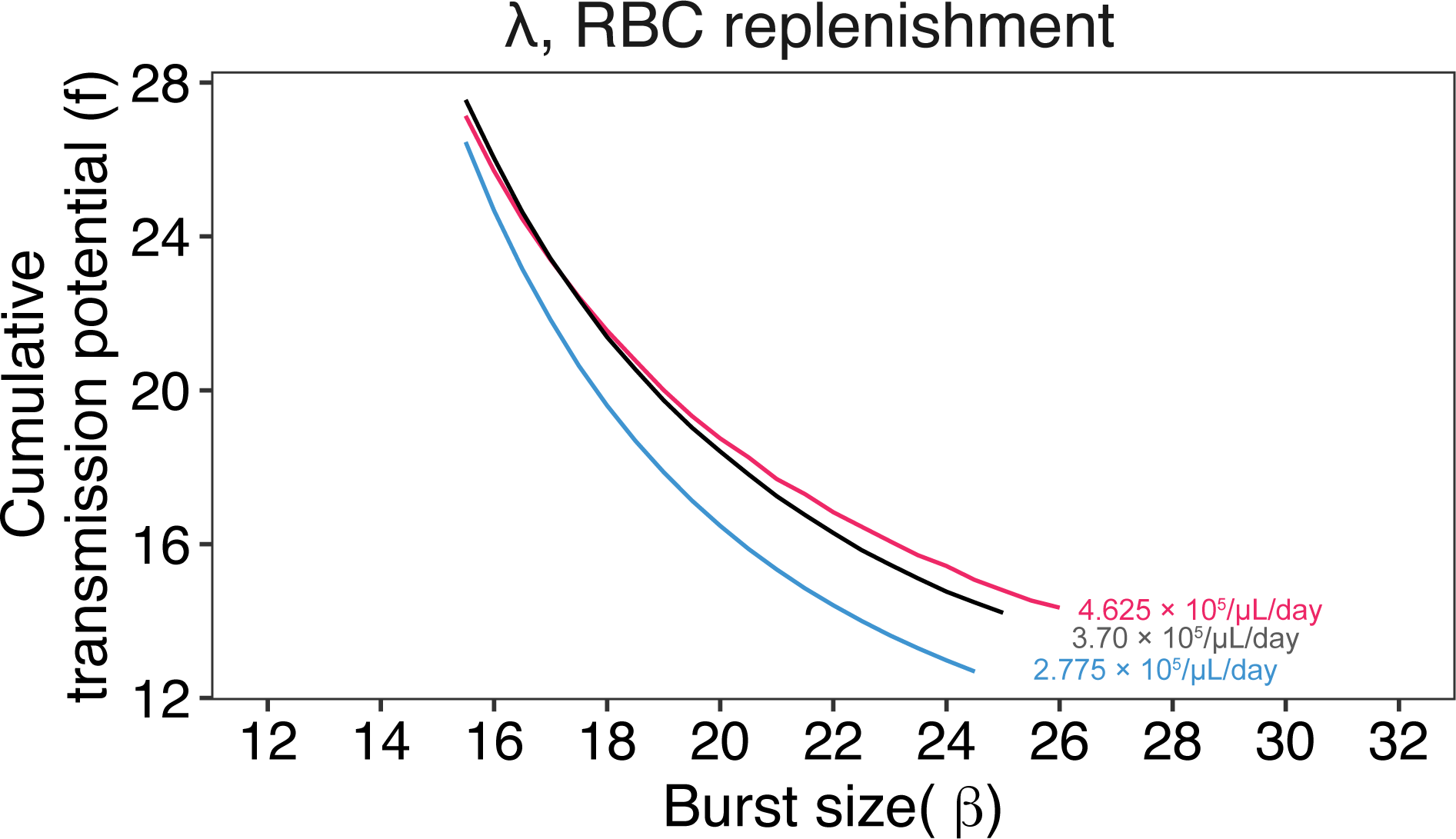
The optimum burst size (*β*) does not change when the replenishment rate of susceptible RBCs (*λ*) is decreased (blue line) or increased (red line) by 25%.

**Figure S9:**
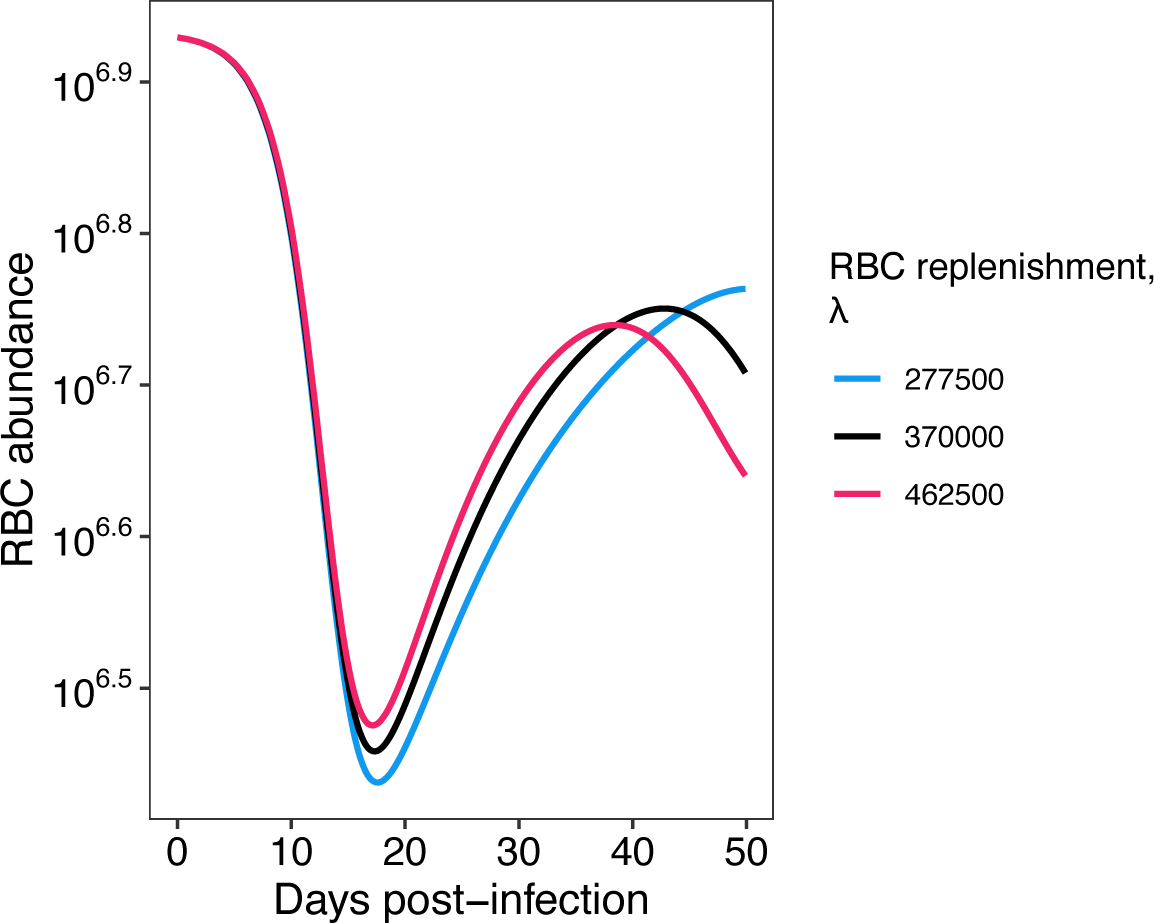
Changes to the maximum RBC replenishment rate (*λ*) do not affect optimal burst size because the effect is only apparent after the peak parasite load. We show that for a burst size (*β*) of 15.5 and a transmission investment (*c*) of 76%, the RBC abundance (log10) are depleted at varying rates depending on the different values of *λ*. The blue and red lines represent the 25% decrease or increase from the original parameter value (black line). Regardless of *λ*, the infections all start similarly, and only when RBCs are depleted is there a difference in the minimum RBCs.

**Figure S10:**
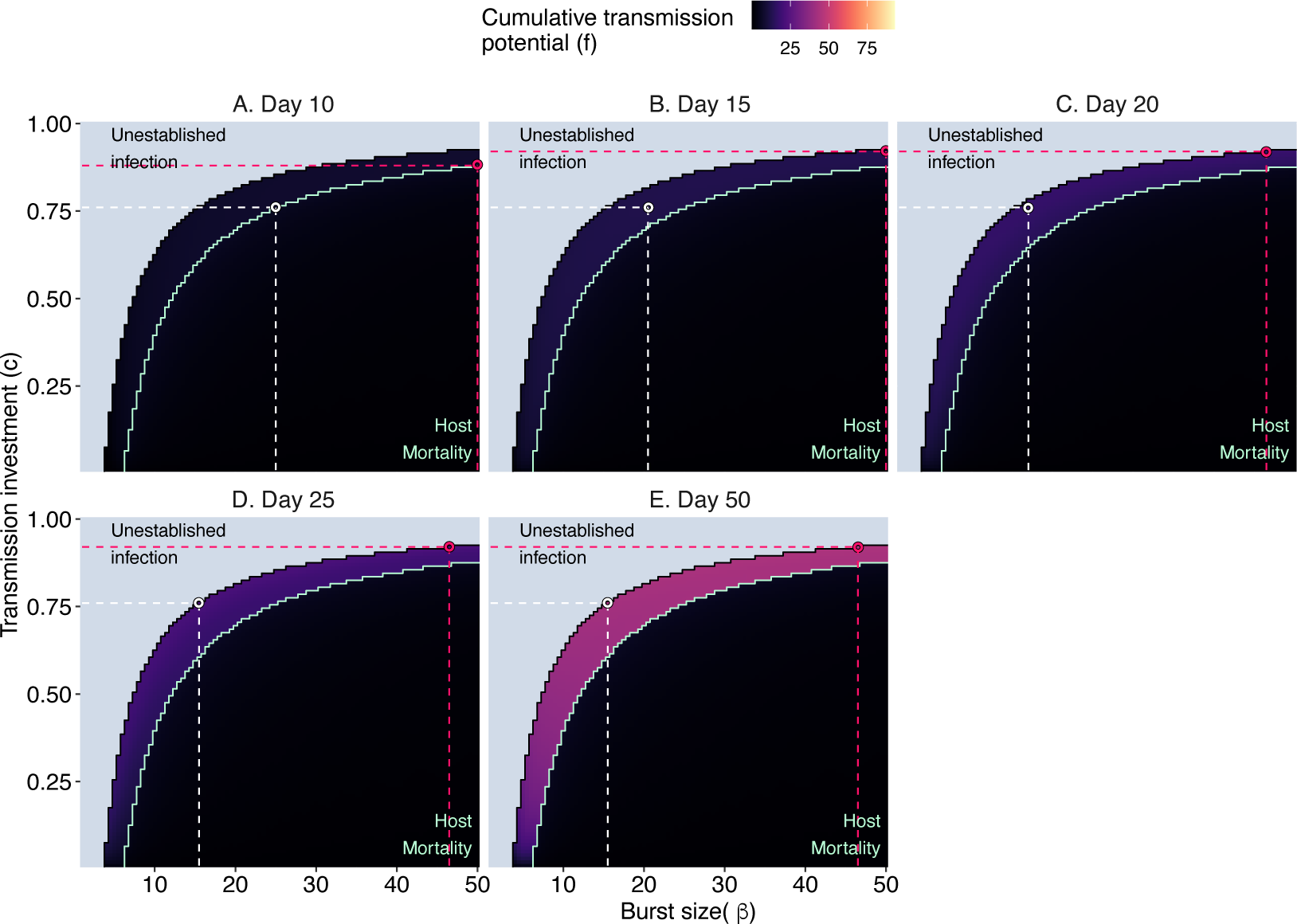
Varying the total infection length, the optimal burst size (*β*) and transmission investment (*c*) shift. The pink points represent the best strategy maximizing cumulative fitness across all parameter combinations. The white points represent the optimal strategy when we fix the transmission investment to 76% (see Figure 6) When the infection lasts (A) 10 days, the optimal strategy (pink points) is *β* = 50 and *c* = 88%. As the infection increases to (B) 15 days, the transmission investment increases and the strategy is *β* =50 and *c* = 92%. If the total infection lasts between 20 to 50 days (C-F), the optimal strategy is *β* = 46.5 and *c* = 92%. The area in blue at the left and top indicates parameter combinations that do not establish infections (*R_M_* (0) < 1.5) and the cumulative transmission potential (*f*) is set to 0. The green line indicates the boundary where the *β* and *c* induce host mortality due to catastrophic loss of host RBCs.

**Figure S11:**
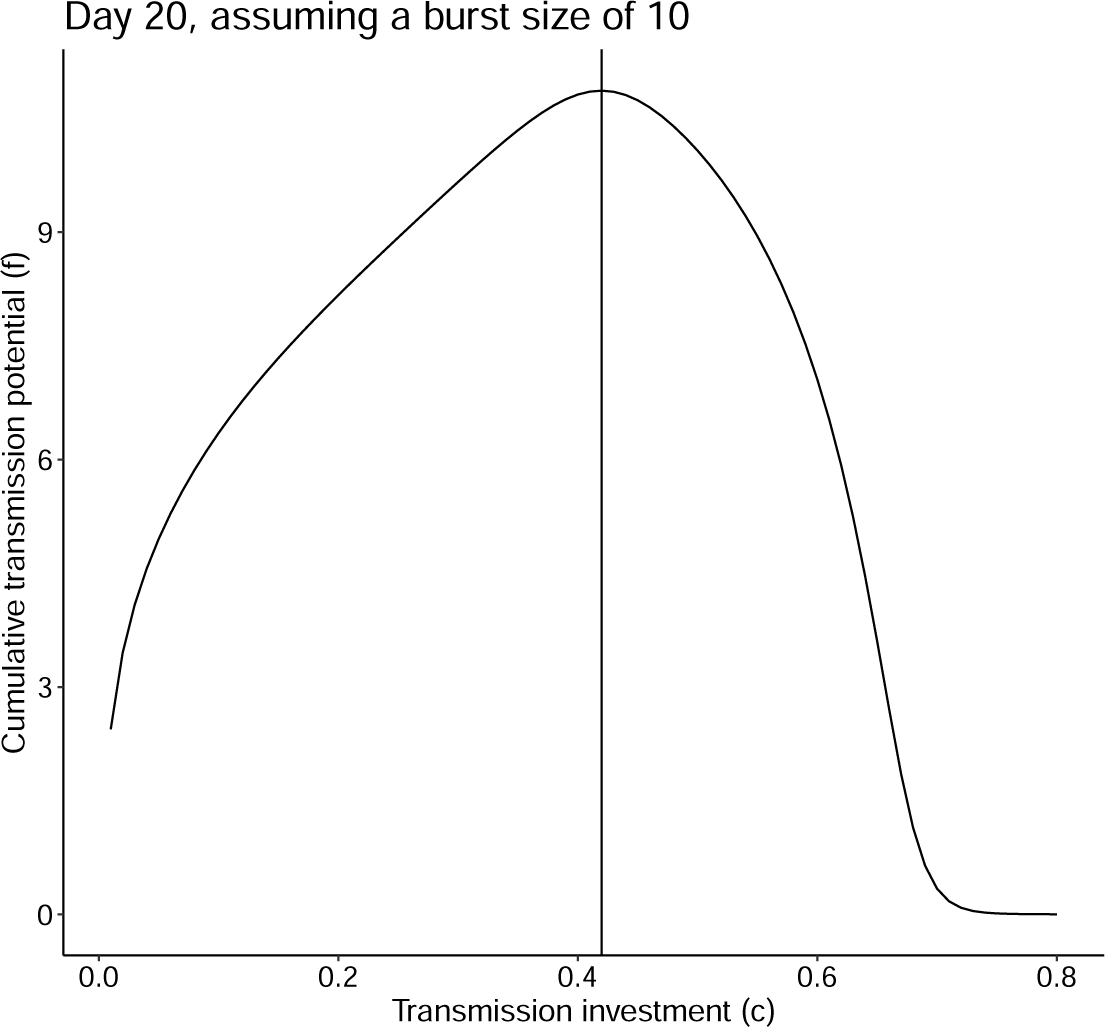
Fixing the burst size (*β*) to 10 (as in Greischar *et al*., 2016), we likewise find that the optimal transmission investment is 42% for a 20 day infection assuming there is no host mortality. This optimal transmission investment remains unchanged even when we assume mortality.

